# Lipid Nanoparticles for Spleen-Targeted RNA delivery

**DOI:** 10.64898/2026.04.13.718229

**Authors:** Kseniia Vlasova, Noorjahan Aibani, Mrinmoy Sanyal, Marco Herrera, Avisek Deyati, Ekram Helmy, Harvie Pierrot, Seba Jumaa, Daniel Arriaza, Miao-Chih Tsai, Ravindra Majeti, William J. Greenleaf, Anitha Thomas, Howard Y. Chang

## Abstract

Lipid nanoparticles (LNPs) formulated with neutral helper lipids efficiently deliver RNA to the liver in pre-clinical models and humans but achieving clinically relevant delivery to other tissues remains a major challenge. To reduce liver uptake, targeting strategies often range from active targeting relying on antibodies to quasi-active targeting by employing permanently charged helper lipids which influence biodistribution after administration. In this study, we present an alternative approach based on varying ionizable lipids and stabilizers, along with optimizing formulation parameters for targeted delivery of circular RNA via a passive targeting approach. We generated a library of 216 LNP formulations and evaluated their performance *in vitro* in Jurkat cells and human primary T cells. The lead LNPs showcasing activity in both Jurkat and T cells were explored for their efficacy *in vivo* via multiple routes of administrations. Our results show that both the identity of stabilizer and ionizable lipid had effects on decreasing hepatic vs. splenic delivery while enhancing splenic accumulation. In line with this improved tissue tropism, spleen-tropic LNPs induced distinct transcriptomic remodeling *in vivo* compared with conventional, FDA-approved SM-102 LNPs. These findings demonstrate that extrahepatic targeting of LNPs can be achieved without altering charge of the LNPs and further reveal that hepatic de-targeting efficiency could be influenced by the immune status of the recipient.

## Main

Lipid nanoparticles (LNPs) are FDA-approved for non-viral delivery of nucleic acid-based therapeutics [1,2]. LNPs are currently in advanced clinical testing for delivering diverse DNA and RNA-based therapeutic modalities around the world in areas including but not limited to gene therapies, vaccines, cancer immunotherapies, and gene editing. The primary factor limiting broader clinical applications of LNPs is their inherent affinity for the liver due to LDL receptor-ApoE interaction in hepatocytes [3]. Extrahepatic tissue- and cell-specific targeting of LNPs has been achieved not only through the synthesis of new ionizable lipids or active targeting using peptides and antibodies, but also by introducing additional lipid components or by optimizing the ratios of existing components, which influence particle circulation time and biodistribution *in vivo* [4,5]. The common LNP screening strategy typically involves performing *in vitro* testing of a particle library and then advancing the best-performing candidates to *in vivo* studies. However, surprising little correlation is observed between *in vitro* and *in vivo* screening results [6], indicating that successful formulation optimization through *in vitro* testing requires careful selection of appropriate cell lines and signaling readouts to achieve reliable data. In addition, RNA-loaded LNPs have been shown to be highly immunomodulatory, and retargeting from the liver to the spleen may potentially result in increased delivery to effector cells which could lead to unintended immune responses and toxicity [7, 8]. To avoid these potential effects, we aimed at achieving spleen targeting without surface functionalization, or charged helper lipids, by changing the lipid components for circular RNA (circRNA) delivery and subsequently evaluating the effects of these particles on the immune system *in vivo*. To accomplish this, we conducted *in vitro* screening to identify lead LNP formulations and then studied their behavior *in vivo*. We used circRNA as the cargo because it is an increasingly attractive RNA platform with the potential for improved stability and durability [9].

### Lipid chemistry and molar ratio affect tropism of circRNA expression in T Cells

In this study, we systematically varied the chemistry and compositions of 10 Cytiva proprietary ionizable lipids and 3 stabilizing lipids including PEG(2000)-DMG along with helper lipid (DSPC) and a sterol, cholesterol, generating a library of 216 formulations along with preset control LNPs. Across these, we screened lipid molar ratios in the following ranges: 15 - 45% ionizable lipid, 0 - 60% DSPC helper lipid, 0 - 60% Cholesterol, and 0 - 4% stabilizing lipid (Figure 1a). Key formulations within our library were prepared in replicate (Figure 1b-c). Crucial physicochemical properties of the LNPs - including particle size, PDI, and encapsulation efficiency (EE%), along with luciferase protein expression in immune cells were evaluated to identify optimal formulations for *in vivo* studies. From the splenic immune cell populations, we selected Jurkat cells (an immortalized T lymphocyte cell line) and human primary T cells for in vitro studies, as they are widely regarded as difficult cell types to transfect and are highly relevant therapeutic targets for autoimmune, cancer and gene/cell therapy applications.

**Figure 1.**
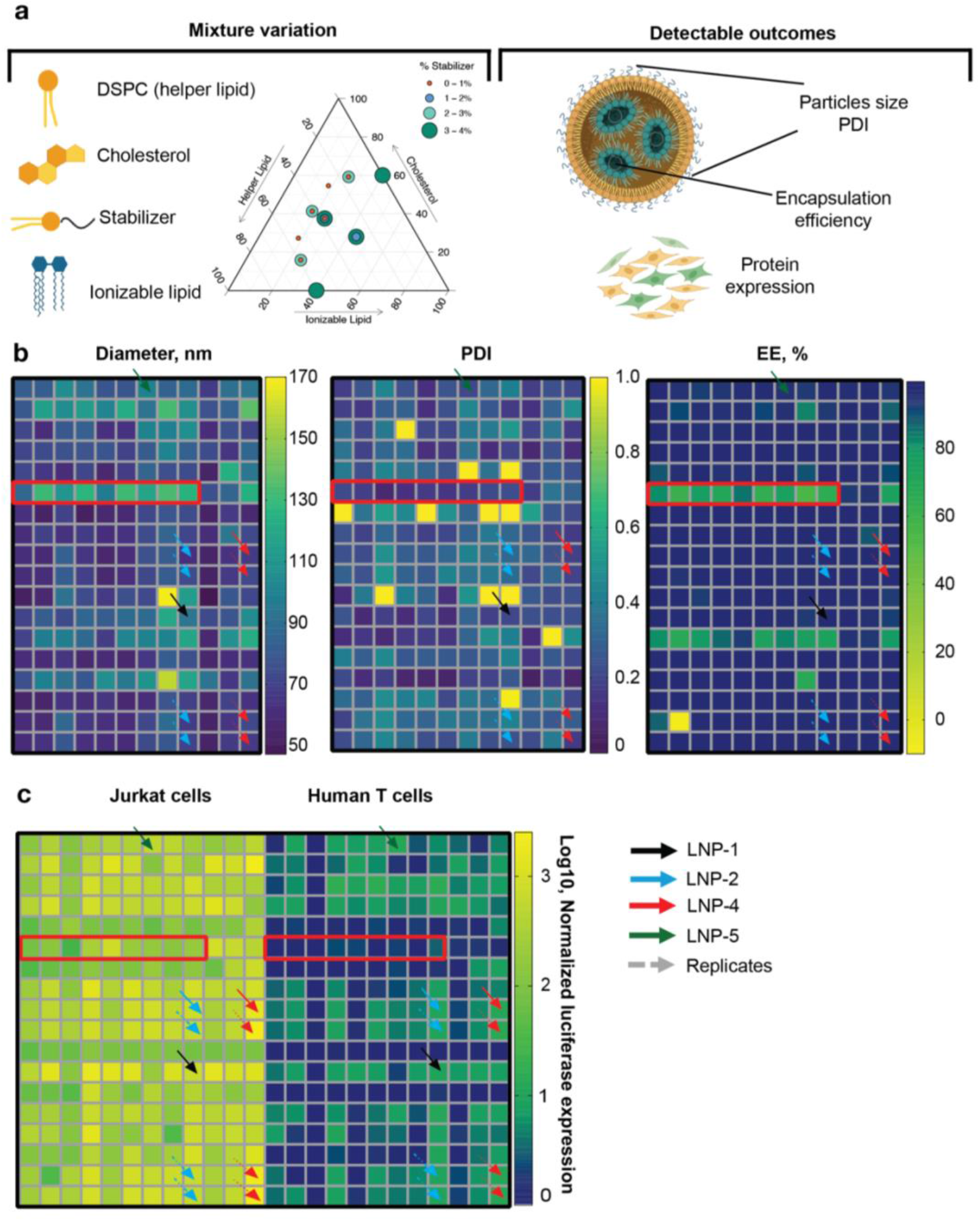
In vitro screening of spleen targeted LNPs. **a)** scheme of the experiment of library preparation and main outcomes. The mages were created with BioRender.com. **b)** physical-chemical characteristics of the produced library. **c)** in vitro luciferase expression in Jurkat cells and human T cells 24 hr post treatment with Fluc circRNA at dose 100 ng per well. Signal was normalized to positive control Cytiva T cells kit. Data are presented as Mean (2 biological replicates with 2 technical replicates each).

Heatmaps of the physical parameters of the particles (Figure 1b, Supporting information Table S1 a-c) showed about 85% of LNPs with particle sizes under 100 nm and encapsulation efficiency (%EE) greater than 85%. Owing to the novelty of the formulation parameters, some formulations exhibited large particle sizes and encapsulation efficiencies (%EE) significantly below the usable average. These formulations contain more than 50% of DSPC and/or cholesterol in the composition, consistent with the previous reports on the effect of high helper lipid and sterol ratios on LNP structure, size, activity and stability [10]. As expected, this group of larger particles did not show detectable functional activity as evidenced by poor protein expression in cells following endocytic escape (Figure 1c, red frame). To note, transfection efficiency in Jurkat cells correlated closely with that observed in primary human T cells (Figure 1c).

While 85% of LNPs had size, PDI, and %EE within our threshold, a substantial number of the formulated LNPs did not achieve our pre-determined threshold level of luciferase expression for further testing (Figure 1). Low transfection efficiency could also be due in part to inherent difficulty of transfection in primary human T cells (Figure 1c), albeit to a lower degree. We also varied the stabilizer lipid such as DMG-PEG2000 molar ratios within the LNPs composition because it was shown previously that increase of stabilizer in lipid compositions significantly decreases the protein expression [11, 12]. We suggest this reduction is due to reduced cellular uptake, endosomal escape, as well as an ultra-stable encapsulated state not conducive to cytosolic delivery. Among the ionizable lipids tested, we observed that higher protein expression was associated with ionizable lipids with pKa ranging from 5.8-6.3. For comparison, the pKa of the SM102 lipid – commonly used for intramuscular delivery of LNPs and known for its high liver tropism after intravenous injection – is 6.7 [13].

We identified five top-performing novel LNP formulations four of which came from the DOE design (LNP-1, LNP-2, LNP-4 and LNP-5) and one from preset Cytiva control formulations added along with the DOE (LNP-3, C11-Supporting information Table S1) to validate further in *in vivo* models. We selected formulations demonstrating human T-cell transfection containing different ionizable lipids and stabilizers to evaluate their *in vivo* behavior. LNPs that had a low stabilizer concentration were excluded from *in vivo* considerations even though they exhibited high activity in cell screen. Transitioning from high-throughput formulation to standard bench scale manufacturing (NanoAssemblr™ Ignite) of the selected LNPs resulted in modest changes in particle physicochemical parameters (Figure 1b, Supporting information Table S1 and Figure 2a). The top performing LNPs overall particle size ranged at or under 120 nm in size and %EE greater than 85%. Next, we checked if high luciferase expression in primary T cells and Jurkat cells were due to immune cells tropism or general improvement of LNPs transfection efficiency. To address this question, we tested top LNPs (Figure 2a) in other, non-immune cell lines such as HepG2 (liver hepatocytes), A459 (lungs epithelium), HEK 293T (kidney epithelium), C2C12 (muscle) cells compared to Jurkat cells (Figure 2b). It was previously shown that some LNPs with immune cell tropism could demonstrate lower transfection efficiency in other cell types like the liver-derived cell line when transfected at equimolar amounts [12]. Potentially there could be a slight difference in LNP association with adherent cells (in our case, HepG2, HEK293T, A549, C2C12) and with suspension Jurkat cells [14]. Recent work by *Liu et al*. showed that SM102 particles had comparable uptake level by Jurkat and HEK293T cells, but different cytosolic mRNA delivery and protein expression [15], which suggests that cell tropism is not exclusively identified by only cellular uptake. Our standard SM102 control demonstrated higher luciferase expression in muscle cells compared to all other cell lines (Figure 2b), which is in line with Moderna SM102 original formulation’s target [16]. In contrast, out of our top-hit selection, we did not detect strong cell-specific tropism patterns (Figure 2b), however, LNP-3 and LNP-4 showed higher tropism to Jurkat and HepG2 cells compared to A549 or C2C12 cells. LNP-5 exhibited broader transfection efficacy across cell lines with smaller difference in efficiency, but highest transfection of Jurkat and HepG2 cells.

**Figure 2.**
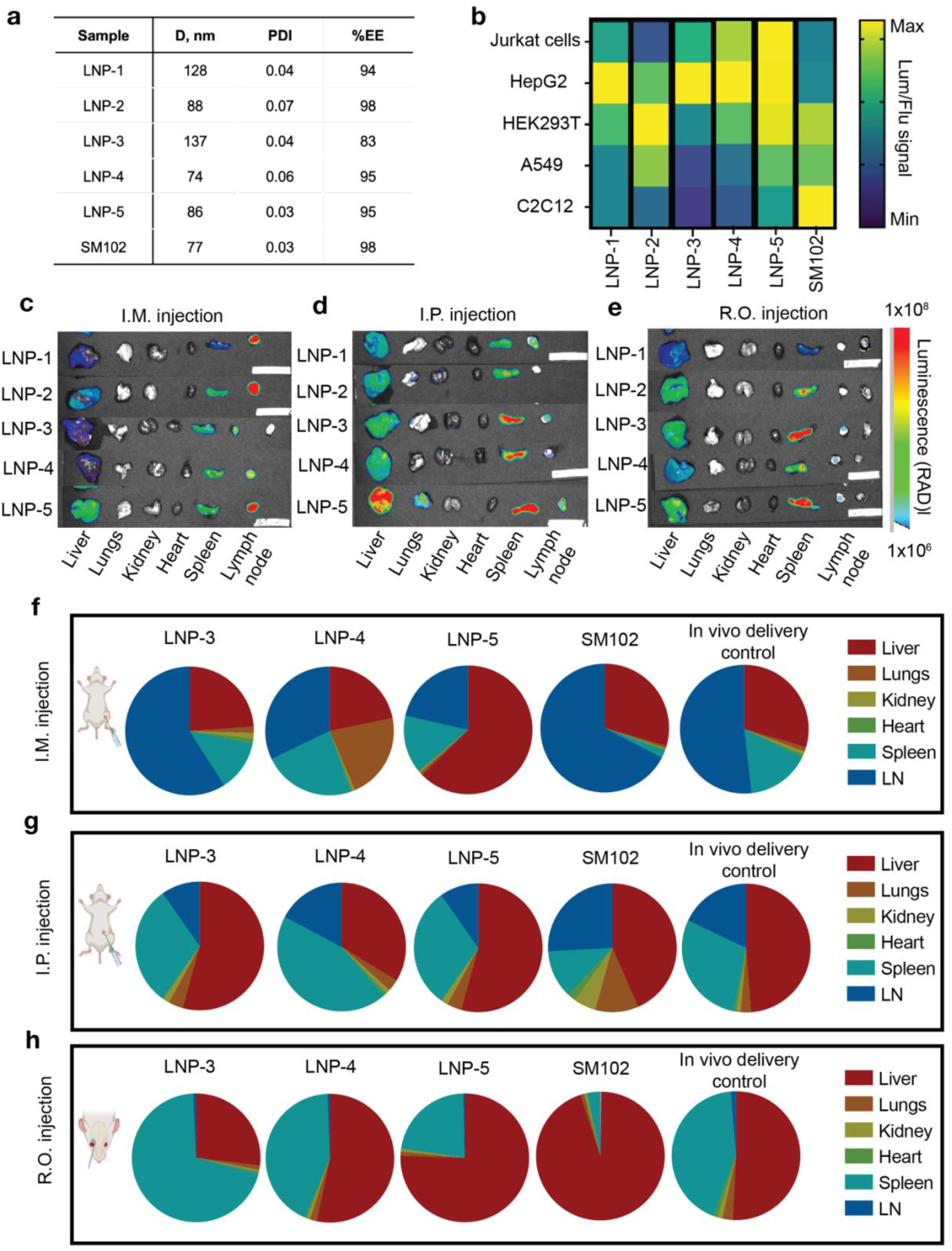
In vivo evaluation of top spleen targeted LNPs. **a)** physical-chemical characteristics of the LNPs. **b)** in vitro luciferase expression across cell lines 24 hr post treatment with Fluc circRNA at dose 100 ng per well. Luminescence signal was normalized to fluorescent signal from live cells. Each column (LNP type) was normalized to PBS control separately. Data are presented as Mean (2 biological replicates with 6 technical replicates each). **c-e)** representative ex vivo images and **f-h)** biodistribution estimations of luciferase expression in organs 6 hr post i.m., i.p. and r.o. administration of LNPs with Fluc circRNA at dose 2 μg per BALB/c mouse. 3-5 biological replicates, 2 batches of each LNPs type. Mice images were created with BioRender.com

Deng X et.al. reported that LNPs with a pKa of 4.7-5.2 are more efficiently delivered to the spleen due to their surface charge at neutral pH [17]. For LNP screens in immune cells we considered pKa of the ionizable lipid as the primary factor for influencing cell transfection efficiency and used ionizable lipids with pKa range of 5.8 -6.3. In addition, the actual lipid ratios in the formulation - particularly the amounts of the helper lipid (DSPC) and the sterol (cholesterol) - also play a significant role. Recently, a new generation of LNPs with a more liposome-like structure showed extrahepatic accumulation *in vivo* by increasing the molar ratio of helper lipid and cholesterol relative to the ionizable lipid [10]. *Cheng et al*. demonstrated, using the MC3 ionizable lipid, that particle stability and delivery efficiency strongly depend on a maximal threshold of helper lipid and cholesterol content, with the optimal level being lower 40% for each component [10]. For our ionizable lipids, we found that the optimal amounts of DSPC and cholesterol are 50% and lower for each. For the third component varied, the stabilizer, several strategies have been reported to increase extrahepatic delivery of LNPs, including the use of PEG-lipids with longer acyl chains (e.g., C18 instead of C14), PEG alternatives such as polyoxazolines and polysarcosines, and higher proportions of these in the formulation [18, 19, 20]. In our *in vitro* screening, the type of stabilizer did not significantly affect performance; however, lower amounts of stabilizer were associated with higher protein expression, possibly owing to a balance between particle stabilization and delivery capacity [11]. When we evaluated our top LNPs in non-immune cell lines, we found that LNP-3, LNP-4, and LNP-5 exhibited higher expression in Jurkat cells and HepG2, compared to epithelium HEK293 T or muscle cells. Such type of tropism is quite uncommon and interesting and may be due to different ionizable lipids and stabilizers present in these LNPs. *Liu et al*. proposed an interesting concept based on similar observations but in different cell types [15]. They tested *in vitro* standard SM102, DLin-MC3-DMA, or ALC-0315 with *in vivo* liver expression across various cell lines and showed that 1) protein expression depended on other factors beyond endosomal escape, and possibly on the intracellular stability of the different LNP formulations and the rate at which the different formulations release mRNA in the cytosol; 2) for the same formulations endosomal escape efficiency differed across the cell types. We would add that the rates of uptake and the cell-cycle state of specific lines also can contribute to the differences in reporter expression.

### In vitro screening correlates with in vivo results and top LNPs deliver to extrahepatic tissues

We next assessed whether the *in vitro* screening results correlated with *in vivo* biodistribution. We tested the top formulations (Figure 2a) *in vivo* using different routes of administration and used the *in vivo* RNA delivery kit with ionizable lipid from the same family as our screening library and SM102 LNPs as control LNPs. After intramuscular (i.m.) injection, our top LNPs showed higher spleen tropism compared to the standard SM102 formulation (Figure 2c and 2f). In addition, some of these formulations drained to the inguinal lymph node adjacent to the injection site (Figure 2c), as reported previously [21]. Notably, expression kinetics for LNP-3, LNP-4 and LNP-5 from muscles were lower compared to SM102 formulation at the same dose, but signal from spleen was the same or higher (Figure 2f, Supporting information Figure S1). A comparison of the *in vitro* transfection results with the *in vivo* biodistribution after i.m. administration showed similar patterns. Intraperitoneal (i.p.) and retro orbital (r.o.) administration also resulted in substantial accumulation of these LNPs in the spleen, accompanied by reduced liver uptake relative to control formulations (Figure 2d-e, 2g-h). As reported earlier we also observed that SM102 LNPs have higher propensity to go to liver, and the extent of translation was dependent on the route of administration [13, 17]. The *in vivo* delivery control exhibited organ distribution profiles similar to our top LNPs, and SM102 LNPs were used as the reference for liver-targeted control for subsequent analyses to streamline comparisons. R.o. injection revealed some correlation for LNP-3, LNP-4 and LNP-5 with in vitro results. Interestingly, LNP-5, which showed high protein expression in epithelial cell lines (HEK293T and A549) with a shift of transfection efficiency toward Jurkat and HepG2 cells (Figure 2b), exhibited higher tropism to the liver than to the spleen *in vivo* (Figure 2h). Similarly, the SM102 formulation, which also demonstrated higher *in vitro* protein expression in epithelial cells (HEK293T and A549) compared to Jurkat or HepG2 cells (Figure 2b), showed strong liver tropism *in vivo*, consistent with previous reports [22]. Flow cytometry analysis confirmed that LNP-3, LNP-4 and LNP-5 transfect immune cells in spleen (macrophages, T cells and B cells) (Supporting information Figure S2) following *in vivo* administration. At the same time, the SM102 formulation also showed luciferase expression in immune cells (Supporting information Figure S2). Interestingly, we observed an increase in spleen signal for LNP-3 and SM102 with dose escalation (Figure 3a-b), while the liver signal remained unchanged (Supplementary Figure S3). LNP-4 and LNP-5 at 2 µg dose of Fluc circRNA showed approximately the same signal in both the liver and spleen as a 4 µg dose (Supporting information Figure S3). We observed our top spleen-targeted LNPs accumulated in the liver, but showed a significant shift toward spleen protein expression, consistent with the *in vitro* screening results across multiple cell lines. We note that because liver-resident immune cells contribute to the overall luciferase signal detected in the organ, an approach that incorporates *in vitro* screening of LNP libraries using multiple cell lines can help optimize and select formulations with more efficient organ targeting *in vivo*. This could be relevant for circRNA payloads, where IRES-driven translation may make expression outcomes highly sensitive to formulation-dependent cytosolic release [23, 24].

**Figure 3.**
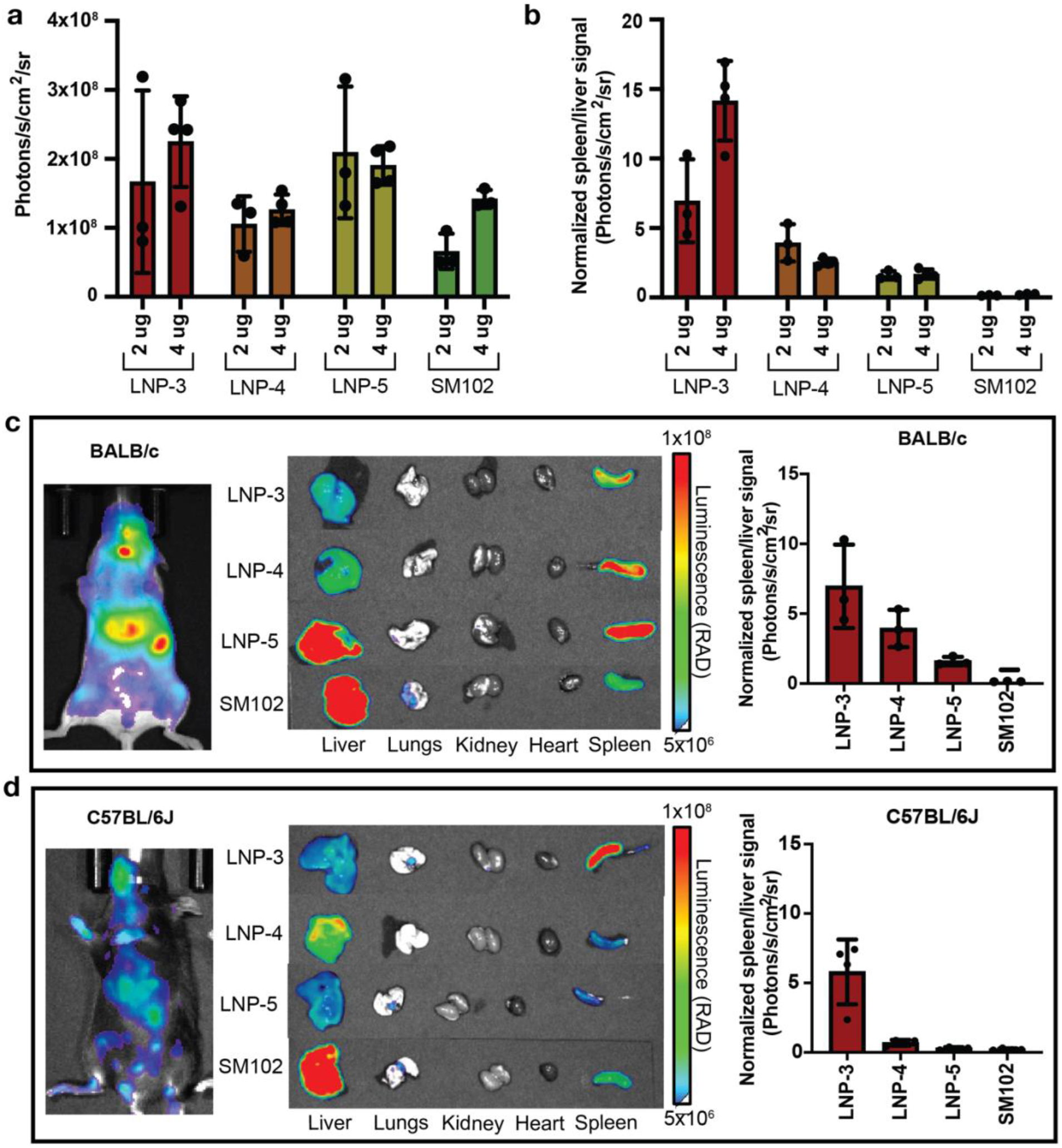
Comparison of spleen tropism of LNPs between mice strains. **a)** dose dependence of bioluminescent signal in spleen (BALB/c mice) and **b)** normalized spleen to liver signal 6 hr post r.o. injection of LNPs. Data are presented as Mean±SD (n=3-4 biological replicates). **c-d)** ex vivo biodistribution of LNPs and spleen to liver signal in **c)** BALB/c mice and **d)** C57BL/6j mice 6 hr post r.o. injection of 2 μg of Fluc circRNA. Data are presented as Mean±SD (n=3 for BALB/c mice and n=4 for C57BL/6j mice).

### Spleen-targeted LNPs potentially activate Th1 and Th2 pathway

Recent studies on pharmacokinetics of LNPs explained the effect of lipid composition on *in vivo* biodistribution and blood clearance frequency by i) protein corona through addition of constantly charged lipid (SORT LNPs) [25] and by ii) upstream or downstream of cellular signaling pathways through variation of neutral helper lipids (glycolipids) [26] or ionizable lipids [27]. Since LNPs are amenable to interaction and subsequent expression in immune cells, we examined whether their biodistribution depends on the basal immune system. We compared luciferase expression after r.o. administration of LNPs in BALB/c and C57BL/6J mice (Figure 3c-d), which are known to exhibit distinct differences in susceptibility to various pathogens [28,29]. On average, we did not detect a significant difference in spleen distribution for LNP-3 or the SM102 formulation, but LNP-4 and LNP-5 showed reduced protein expression in the spleen in C57BL/6J mice. Species-dependent responses to LNPs were previously reported by *Hatit et al* [30]. This work observed differences in translation and endocytosis in hepatocytes across species, which were attributed to differential immune responses contributing to strain-dependent LNP delivery.

Cytokine analysis 6 hr post r.o. injection of LNPs in BALB/c mice showed that LNP-3, with the highest spleen tropism, modulated immune activation, including recruitment of monocytes/macrophages (IL-6, IL-17A; CCL2, CCL7, CCL4), neutrophils (TNF-α, G-CSF; CXCL1), and eosinophils (IL-5), consistent with moderate activation of innate pro-inflammatory pathways and early myeloid-driven immune priming in the spleen (Figure 4a, Supporting information Table S2). Above immune reaction is likely attributable to a combination of formulation parameters used. Other particles exhibiting lower spleen targeting showed milder secretion of cytokines activated monocytes and neutrophils, but significant level of IL-5. Cytokine profiles of LNPs are found to be distinct likely due to compositional differences. All LNPs tested *in vivo* showed no signs of toxicity from its clinical symptoms during the study.

**Figure 4.**
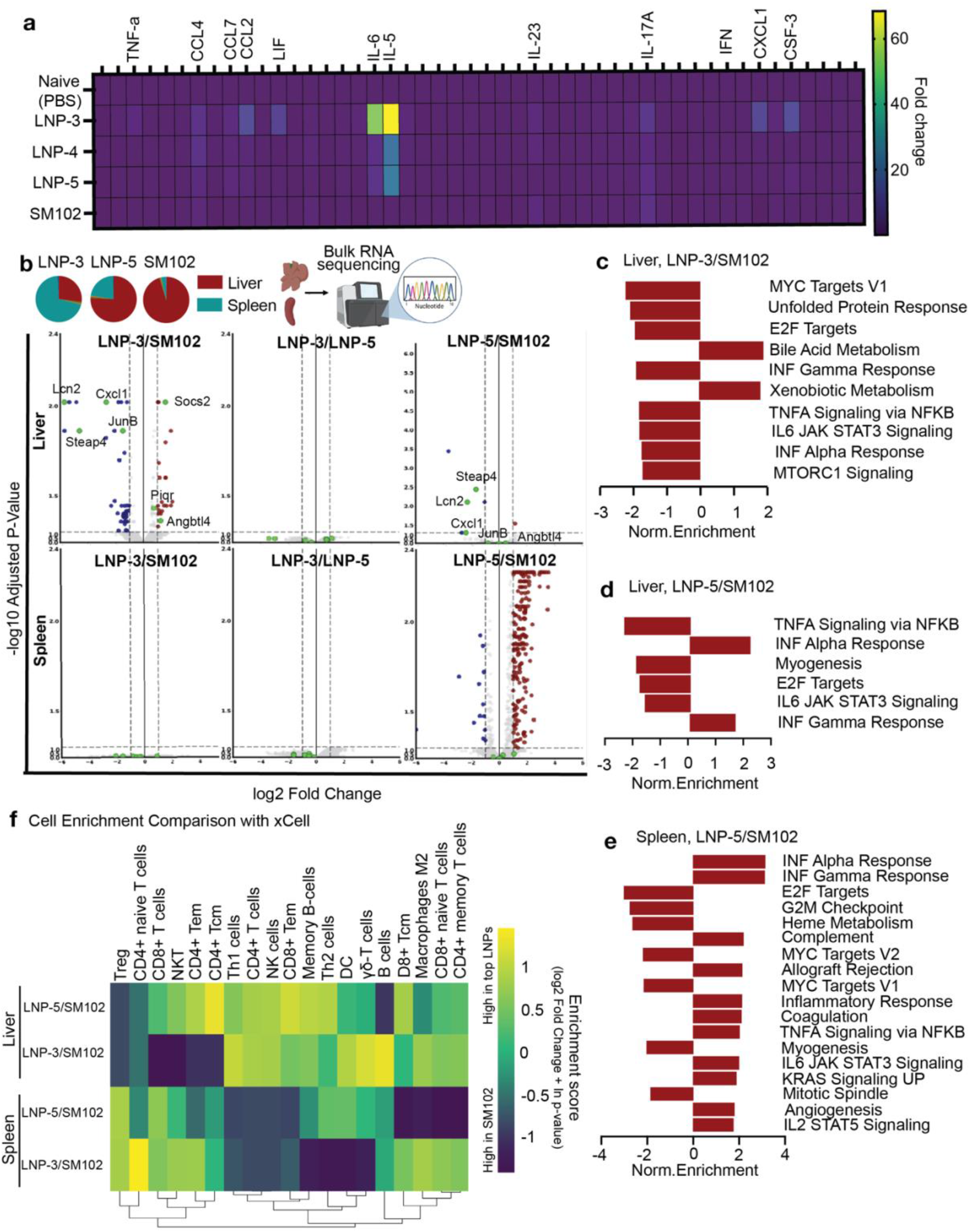
Impact of spleen-targeted LNPs on in vivo immune response. **a)** cytokines screening in serum from BALB/c mice LNP-3, LNP-4 and LNP-5 produced more IL-5 and IL-6 than SM102 at 6 hr post administration. Fluc circRNA dose 2 μg. Data are presented as Mean (n=3 biological replicates). Data are normalized to naïve mice. **b)** RNA sequencing of samples from liver and spleen. Volcano-graphs of up-regulated and down-regulated genes in comparison groups (LNP-3 to SM102, LNP-3 to LNP-5, LNP-5 to SM102). The image was created with BioRender.com. **c-e)** pathways enrichment in groups of comparison. **f)** cell enrichment analysis in groups of comparison with xCell. BALB/c mice, 6 hr post administration. Fluc circRNA dose 2 μg. n=3 biological replicates per group.

We performed bulk RNA-sequencing on liver and spleen, 6 hr post r.o. administration of particles with highest spleen tropism (LNP-3), particles with spleen and liver tropism (LNP-5) and SM102 control (Figure 4b). Our analysis revealed that 40 genes (including JunB, Steap4, Cxcl1) in liver were down-regulated and 21 genes (including Socs2, Piqr) up-regulated after LNP-3 treatment compared to SM102 formulation. The most strongly enriched pathways in the LNP-3 group point to metabolic processes compared to SM102 (bile acid metabolism and xenobiotic metabolism) in liver; LNP-3 demonstrated reduction in inflammatory signaling and cellular stress and proliferation in liver (Figure 4c). Importantly, LNP3 did not induce any significant differences in transcriptomic activity that were detected in spleen (Figure 4b).

When looking at LNP-5’s effect on gene expression, we detected down-regulation of several genes including Saa2, Saa3, Saa4, Cxcl1 in liver, suggesting a global down-regulation of core pro-inflammatory signaling and proliferative and stress markers (Figure 4d). At the same time LNP-5 caused induction of a high-interferon environment (INF alpha response and INF gamma response) unlike to LNP-3 case (Figure 4d). Contrary to LNP-3, LNP-5 showed significant up-regulation of genes in spleen compared to SM102 formulation. These particles mainly activated

INF alpha and INF gamma pathways and down-regulated E2F targets, G2M checkpoints and heme metabolism. Comparison of LNP-3 and LNP-5 in liver and spleen did not reveal significant gene expression changes.

Next, we analyzed cell enrichment in groups compared to SM102 formulation using xCell [31]. In the context of bulk RNA-seq, enrichment of cell-type-specific gene signatures provides the information whether a particular cell population is expanded or depleted. xCell analysis enables us to clarify if our difference in gene expression between top LNPs and SM102 formulation is due to a change in the gene’s expression or in the cellular compositions between the conditions [32]. Overall, we did not observe a drastic change in cellular compositions in liver or spleen after administration of our top LNPs compared to SM102 (Figure 4f). However, we observed few trends based on xCell analysis. These trends are that r.o. injection of both LNP-3 and LNP-5 increased regulatory T and CD8+ T cells in the spleen relative to SM102. This may indicate that LNP-3 and LNP-5 potentially can induce a more immunosuppressive or tolerogenic microenvironment in the spleen [33]. In line with this, LNP-3 administration resulted in a higher number of CD4+ naïve cells and lower DCs in the spleen compared to SM102, which could be a result of Treg-mediated suppression [34]. We also observed enrichment of B cells and γδ-T cells in the liver (Figure 4f) after LNP-3 administration compared to SM102.

Thus, RNA sequencing data suggested that LNP-3 acted as a low-inflammatory, particle in the liver. In contrast, the formulation of LNP-5 acts as an immunostimulant, showing a distinct ability to significantly upregulate interferon pathways in both the liver and spleen. Such up-regulation of genes compared to SM102 control indicates that LNP-5 with partial spleen tropism activates Th1 innate immune pathway. These results are further complemented by the cytokines screening data (Figure 4a). Notably, LNP-3 administration led to higher IL-5 expression than LNP-5. Taken together, the finding that LNP-3 reduced the Th1/pro-inflammatory pathway (compared to SM102) and increased IL-5 cytokine levels likely indicates a shift toward Th2 polarization. When compared, our *in vivo* results from two strains (Figure 3c-d) LNP-4 and LNP-5, which induced moderate IL-5 activation, showed higher signal in BALB/c mice than in C57BL/6J, while LNP-3 exhibited constant spleen to liver signal ratios across strains. C57BL/6J mice are Th1-dominant, leading to robust, inflammatory responses with strong neutrophil/M1 macrophage recruitment, while BALB/c mice are Th2-dominant, thus favoring antibody responses (humoral immunity) with slower, more M2 macrophage-driven response [28, 29]. LNP-3’s higher spleen tropism may be attributed to recruitment of the Th2 immune arm (monocytes, neutrophils, and eosinophils) and the significant dampening of the Th1/interferon/TNF response.

Our findings correlate with recently published data that spleen-target LNPs induced inflammation and increase of cytokines in the serum [26]. Given that the administered LNPs were well tolerated at the efficacious doses tested and did not induce observable clinical symptoms, it is reasonable to assume that spleen-targeted LNP delivery – a lymphoid organ – may stimulate resident immune cell populations, leading to modulation of cytokine profiles. Such immune modulation is likely to be species-dependent for a given LNP formulation, and distinct LNPs may induce different immunomodulatory responses. While this may be beneficial for vaccine applications, for other therapeutic approaches such as gene editing or gene delivery in general, the addition of inflammation suppressors could be a potential solution. Finally, as previous research pointed out [26, 30], our study was also conducted in mice; in other species, spleen tropism as well as the immunogenicity profile may differ. Our trends and relationships should be interpreted within this formulations design landscape and future work across species biodistribution may be necessary to determine the extent of which these rules apply universally. The temporal dynamics of this immune modulation, including onset, duration, and potential reversibility, warrant further detailed investigation. Understanding these effects will be particularly relevant for the development of in vivo CAR therapeutics, and the broader implications of LNP-induced immune modulation should be explored further.

Taken together, our results show that spleen-targeted LNPs can be generated and optimized through variation of the ionizable lipid and stabilizer and their formulated molar ratios. This is an alternative approach compared to charge-based SORT technology and helper glyco-lipid based strategy to direct LNPs towards spleen. Liver protein expression can be reduced by combining formulation optimization with cargo optimization, for example, by varying the IRES in the circular RNA construct or by UTR optimization for mRNA. An important consideration for the clinical application of spleen-targeting LNPs is a thorough understanding of their immunogenicity profile – including onset, duration, and potential reversibility – and how this informs potential implications and opportunities for therapeutic versus vaccine applications.

## Author Contributions

H.Y.C., W.J.G., M.C.T., A.T., P.H., K.V., M.S. and N.A. designed research; K.V., M.S., M.H., A.D., E.H., S.J., D.A. and N.A. performed research; K.V., M.S., M.H., A.D. analyzed data; and K.V. wrote the manuscript. All authors reviewed & edited.

## Disclosure

The authors declare the following competing financial interest(s): H.Y.C. is an employee and stockholder of Amgen, is a co-founder of Accent Therapeutics, Boundless Bio, Cartography Biosciences, and Orbital Therapeutics, and was an advisor to 10x Genomics, Arsenal Bio, Chroma Medicine, Exai Bio and Vida Ventures. W.J.G. is a consultant and/or equity holder for 10x Genomics, Guardant Health, Nvidia, Diffuse Bio, Lamar Health and Ultima Genomics and co-founder of Protillion Biosciences. Anitha Thomas, Pierrot Harvie, Avisek Deyati and Noorjahan Aibani are employees of Cytiva. The rest of authors declare no competing financial interest.

## Acknowledgments

This work was generously supported by Cytiva. H.Y.C. and W.J.G. acknowledge support from the Stanford RNA Medicine Program. Cytiva team acknowledges the support of Leung See Wai Sherry for high throughput assay normalization, Wu Tony for ULPC assay, Sharma Ruchi and Anantha Malathi for LNP formulations and Devina Divekar for project management.

