## Supporting information for "Lipid Nanoparticles for Spleen-Targeted RNA delivery"

### Materials and methods

#### Ethical statement

Animals

Female BALB/C, female C57BL/6 (The Jackson Lab) mice, aged 8 weeks, were used in the study. All mice used in our experiments were maintained in the animal facility of Stanford University with a 12 h light cycle and free access to water and food. Mice were provided ad libitum access to standard rodent chow and water. Experimental procedures were conducted during the light phase in accordance with protocol. Blinding was not applied in our studies; we used matched age and sex controls. All experiments were conducted in accordance with the Institutional Animal Care and Use Committee (IACUC, protocol # 14046) of Stanford University.

#### 1 Circular RNA production and QC

Luciferase encoded circRNA (Fluc circRNA) was synthesized in vitro using the T7 RNA polymerase transcription method (Codex® HiCap RNA polymerase, Aldevron, Fargo, ND). circRNA templates were synthesized by cloning DNA fragments into a custom entry vector which contains a T7 RNA polymerase promoter, self-splicing group I introns, and a picornavirus-based IRES. The IVT DNA template (1 µg per reaction) was subsequently degraded with 0.5 µL Turbo-DNAse (NEB, cat #M0303S) for 20 min at 37 °C. The remaining RNA was purified by precipitation with 8M LiCl (Sigma-Aldrich, L7026) and digested with 1U RNase R (made in-house) per microgram of RNA for 60 min at 37 °C to remove linear precursor and spliced-out introns. Samples were then again purified by precipitation with 8M LiCl, quantified using a Nanodrop One spectrophotometer, and verified for complete digestion using an Agilent TapeStation system (Agilent, Santa Clara, CA) or denaturing agarose gel electrophoresis. After dephosphorylation by Antarctic phosphatase (AnP) (NEB, cat #M0289L), RNA was HPLC purified on an Agilent 1260 Infinity II LC System (Agilent, Santa Clara, CA) and using an ion-pair reversed-phase chromatography column (ADS Biotec Inc, Omaha, NE) under thermally denaturing conditions to further enrich for circular RNA and remove possible dsRNA contaminants. Purity and integrity of the finally purified RNA were checked by denaturing agarose gel electrophoresis (Supporting information materials Figure S4). RNA was aliquoted and stored at -80 °C until use.

**2 Formulation of Lipid nanoparticles**

The lipid mixes were made of four different components: Cytiva proprietary ionizable lipids or SM-102 along with a helper lipid, cholesterol and stabilizer. Lipid components dissolved in ethanol were mixed at specific molar ratios totaling at 100 mol%. 2-dioctadecanoyl-sn-glycero-3-phosphoethanolamine (DSPC) and cholesterol were used as the helper lipid and sterol, respectively. SM-102 LNPs were formulated with 50 mol% SM-102, 10 mol% DSPC, 38.5 mol% cholesterol and 1.5 mol% PEG-DMG-2000 (1,2-dimyristoyl-rac-glycero-3-methoxypolyethylene glycol 2000). The Cytiva proprietary ionizable lipids are described in patent application publications such as WO2021000041A1 and WO2020210901A1. The Cytiva proprietary lipid mix composition and the LNPs described in this manuscript can be purchased through Cytiva referring to this publication. The Cytiva RNA Delivery LNP Kit (Cat No: 1002569, 1002471, Cytiva), was used as control for in vivo studies. To formulate the lipid nanoparticles, the circular RNA solution was prepared in a selected formulation buffer at pH 4±0.2. For high throughput screening in Jurkat and primary T cells, circular RNA LNPs were formulated using automated high throughput microfluidic mixing methodology. GenVoy-ILM™ T cell kit (Cat No: 1001144, Cytiva) was used for normalizing the signals as positive control in in-vitro screening (Figure 1c). *In vivo* study LNP formulations were manufactured using the Cytiva NanoAssemblr™ Ignite system with NxGen™ mixing technology. Briefly, the four component lipid mixes at the desired ratios in ethanol were rapidly mixed with circular RNA solution in the formulation buffer at pH 4±0.2 at a Nitrogen-to-Phosphate (N/P) molar ratio of 8, aqueous to ethanol flow ratio of 3:1 and a total flow rate (TFR) of 12 mL $min^{-1}$ in NxGen™ cartridge. All LNPs were subjected to downstream processing. High throughput formulations were dialyzed whereas Ignite formulations were subjected to buffer exchange and concentration using Amicon Centrifugal filter (MWCO 30 kDa) following 25x dilution in 1xPBS (pH 7-7.3, $Mg^{2+}$/$Ca^{2+}$- free) and sterile filtered using 0.22 µm syringe filters. Final LNPs were mixed with Cytiva proprietary cryobuffer in 1:1 ratio and stored at -80°C until utilized.

**3 Characterization of LNPs**

LNPs from high throughput formulation screening were measured via Static Light Scattering (SLS) using the Wyatt DynaPro Plate Reader III. Particle size and PDI of Ignite formulated LNPs were analyzed by Dynamic Light Scattering (DLS using the Malvern NanoZs Zetasizer (Malvern Instruments, UK) in backscattering detection mode (He/Ne laser 633 nm, Measurement angle 173°) and reported as Z-average size. LNPs were prepared in polystyrene cuvettes and appropriately diluted for sizing analysis.

Measurement of encapsulation efficiency (%EE) and RNA concentration was done by a modified Quanti-iT RiboGreen™ assay. Briefly, LNPs were incubated in a 96 well plate at 37°C in 1xTE buffer for 10 minutes in the presence or absence of Triton X-100 (0.6%) to determine free and total RNA, respectively. Ribogreen dye prepared 1:100 in 1xTE buffer was added to each well and the fluorescence intensity was measured at Ex/Em 485 nm/528 nm on Biotek™ Synergy™ H1 Multimode Monochromator microplate reader. A standard curve (0.1-2.5 ug/mL) was prepared similarly to assess the concentration of circular RNA in the LNPs. Encapsulation efficiency (EE) was calculated using the following equation:

EE= 100x {(Total RNA$_{\text{(RNA in Triton TE)}}$ - RNA outside LNP$_{\text{(Ribogreen in TE)}}$)/Total RNA$_{\text{(Ribogreen in Triton TE)}}$}.

4 **In vitro screening**

For high throughput screening of LNPs library in Jurkat clone E6-1cells (#TIB-152, ATCC) and human T cells (were obtained from Stanford Blood Center), $1x10^5$ cells per well were seeded in 96-well plate in RPMI-1640 + 10% FCS + 100 units/mL of penicillin and 100 µg/mL of streptomycin + 1 µg/mL ApoE3. Human T cells were stimulated before seeding with IL-2 (final concentration in media: 100 ng/ml IL-2) and CD3/CD28 beads at 1:1 ratio). Next, samples were diluted with complete media + 1 µg/mL ApoE3 for Jurkat cells and full media + 1 µg/mL ApoE3 + 100 ng/ml IL-2 and added to cells at dose 100 ng of Fluc circRNA.

For screening of top LNPs across Jurkat cells (#TIB-152, ATCC), Hep2G cells (#HB-8065, ATCC), A549 cells (#CRM-CCL-185, ATCC), HEK293T (#CRL-11268, ATCC), C2C12 (#CRL-1772, ATCC), $1x10^4$ cells of each type per well were seeded in 96-well plate in complete media according to ACTT protocols. After 24 hr samples in complete media + 1 µg/mL ApoE3 were added to the cells at dose 100 ng of Fluc circRNA per well.

Samples were incubated with cells 24 hr and then equal volume of BriteLite reagent (Revvity) was added and luminescence signal was measured using plate reader. In the case of screening across cell lines, first, cell viability results (CellTiter-Fluor, Promega) and then luciferase expression data (BriteLite reagent, Revvity) were collected 24 h post-treatment using a plate reader. Luminescent readout (in relative luminescence units) was normalized by cell counts commensurate with fluorescence (relative fluorescence units, RFU).

**5 Bioluminescence imaging**

Fluc circRNA encapsulated in particles at a 2 µg dose was injected via intramuscular (i.m.) or intraperitoneally (i.p.) or retro orbital (r.o.) to female BALB/c mice or C57BL 6J mice. For bioluminescence imaging 6 h post injection, mice received D-luciferin substrate (150 mg/kg) i.p. and were imaged according to the manufacturer's protocol. Image acquisition and analysis were performed using the Lago optical imaging system (Spectral Instruments Imaging) and the manufacturer's software (Aura). Mice were anesthetized by isoflurane inhalation during the procedure. For ex-vivo imaging, mice were sacrificed, different organs were harvested, mocked in 1.5 g/L D-luciferin solution and analyzed.

**6 Cytokine assay from serum**

Blood serum from female BALB/c mice 6 h post injection with 2 µg Fluc circRNA, loaded in LNPs, via r.o. injection was collected and analyzed by The Human Immune Monitoring Center using Mouse Luminex 48plex-HC according to the kit protocol as qualified. The data collected by the instrument software are expressed as Median Fluorescence Intensity (MFI). MFI values for each analyte are collected per individual sample well. Analyte standards, quality controls, and sample MFI values were adjusted for background. Data are presented as relative unites (fold change) normalized to PBS controls.

**7 Bulk RNA sequencing**

Liver and spleen from female BALB/c mice 6 h post injection with 2 µg Fluc circRNA, loaded in LNPs, via r.o. injection was collected and then total RNA was extracted using Direct-zol RNA MiniPrep with Trizol (Zymo research) according to the manufactory's protocol. Extracted RNA were sent to Plasmidsaurus for RNA sequencing and data analysis. Quality of the fastq files was assessed using FastQC v0.12.1. Reads were then quality filtered using fastp v0.24.0 with poly-X tail trimming, 3' quality-based tail trimming, a minimum Phred quality score of 15, and a minimum length requirement of 50 bp. Quality-filtered reads were aligned to the reference genome (M. musculus BALB/cJ) using STAR aligner v2.7.11 with non-canonical splice junction removal and output of unmapped reads, followed by coordinate sorting using samtools v1.22.1. PCR and optical duplicates were removed using UMI-based deduplication with UMIcollapse v1.1.0. Alignment quality metrics, strand specificity, and read distribution across genomic features were assessed using RSeQC v5.0.4 and Qualimap v2.3, with results aggregated into a comprehensive quality control report using MultiQC v1.32. Gene-level expression quantification was performed using feature Counts (subread package v2.1.1) with strand-specific counting, multi-mapping read fractional assignment, exons and three prime UTR as the feature identifiers, and grouped by gene_id. Final gene counts were annotated with gene biotype and other metadata extracted from the reference GTF file (Supporting Information Figure S5).  Sample-sample correlations for sample-sample heatmap and PCA were calculated on normalized counts (TMM, trimmed mean of M-values) using Pearson correlation. Differential expression was done with edgeR v4.0.16 using standard practice including filtering for low-expressed genes with edgeR:filterByExprwith default values. Functional enrichment, when available, is performed using gene set enrichment analysis with gseapy v0.12 using the MSigDB Hallmark gene set.

### 8 Immune cell differential enrichment analysis using xCell algorithm

xCell [1] is a gene-signature-based method that estimates the enrichment of 64 immune and stromal cell types from bulk RNA-seq data [1]. Unlike regression-based deconvolution approaches (e.g., CIBERSORTx), xCell employs a gene set enrichment approach using curated cell-type signatures derived from thousands of purified cell-type expression profiles. The method applies single-sample Gene Set Enrichment Analysis (ssGSEA) followed by a spillover compensation step to reduce cross-contamination between related cell types, producing cell-type enrichment scores that reflect the relative prominence of each cell type in a given sample [2]. The analysis quantifies differences in immune cell composition between formulations using a composite enrichment score that integrates both effect size and statistical significance. The output is a matrix of cell-type enrichment scores (cell types × samples), where higher scores indicate greater enrichment of a given cell type in a sample.

Differential Enrichment Analysis

For each cell type scored by xCell, differential enrichment between LNP formulations was assessed using ordinary least squares (OLS) linear regression within each organ. The model took the form:

$$Score_{i,j} = \beta_0 + \beta_1 \cdot Group_j + \varepsilon_{i,j}$$

where $\mathrm{Score}_{i,j}$ is the xCell enrichment score for cell type $i$ in sample $j$, $\mathrm{Group}_j$ is a binary indicator (Treatment vs. Control), $\beta_0$ is the intercept (mean control score), $\beta_1$ is the treatment effect estimate, and $\varepsilon_{i,j}$ is the residual error.

P-values from the t-test on the treatment coefficient were corrected for multiple testing using the Benjamini–Hochberg (BH) procedure to control the false discovery rate (FDR) across all cell types within each contrast.

Composite Enrichment Score

To produce a single metric that captures both the magnitude and direction of change (effect size) as well as the statistical confidence of that change (significance), a composite enrichment score was defined for each cell type in each contrast. This score integrates the $\log_2$ fold change with the natural logarithm of the p-value:

$$ES_i = \log_2 FC_i + \ln(p_i)$$

where:

- $\boldsymbol{ES_i}$ is the composite enrichment score for cell type $i$
- $\mathbf{log_2}\boldsymbol{FC_i} = \log_2[(\mu_{treatment} + \varepsilon) / (\mu_{control} + \varepsilon)]$ is the $\log_2$ fold change of mean xCell enrichment scores between treatment and control groups, with $\varepsilon = 10^{-9}$ as a pseudocount to prevent division by zero
- $\boldsymbol{ln(p_i)}$ is the natural logarithm of the raw p-value from the linear model t-test for cell type $i$

**9 Flow cytometry**

To study the cell type transfection by LNPs, BALB/c mice were r.o. injected with 4 µg Fluc circRNA loaded into particles. Untreated BALB/c mice were used as a control. 6 hr post-injection mice were sacrificed, spleen was harvested and then dissociated. Single cells were generated with the organ dissociation protocol. Briefly, mouse spleen was minced and filtered through a 70 µm nylon mesh strainer to collect single cells. Cells were washed using flow wash buffer (PBS containing 2 mM EDTA and 0.5% BSA) and collected by centrifugation at 450 × g for 5 min at 4 °C. The pelleted cells were resuspended in 100–200 µl of ACK lysis buffer (BioWhittaker, Lonza) for 5 min at room temperature and quenched with the addition of equal volume of 5 mM EDTA in PBS. After washing, cells were stained with live dead cell staining dye FITC and then incubated with Foxp3 staining buffer (eBioscience) overnight at 4 °C. Cells were further stained with the following fluorochrome-conjugated antibodies: Luc-PE (abcam, #AB237253, 1:100 dilution), B220-BUV605 (BD Horizon, #103244, 1:100 dilution), CD11-Percp5.5 (BD Horizon, #117328, 1:100 dilution), TcrB-APC (BD Horizon, #553174, 1:100 dilution), F4/80-Alexa700 (BD Horizon, #123130, 1:100 dilution). Live cells were stained with DAPI (Invitrogen, #3046150, 1:1000 dilution). For validation, please see the above-mentioned vendors' specification sheets. The Fluorescence Minus One (FMO) control for each sample was cells without Luc-PE staining. The stained cells were analyzed using BD FACSymphony A5.

**10 Statistical analysis**

Unless described otherwise, statistical analyses were conducted using GraphPad Prism.

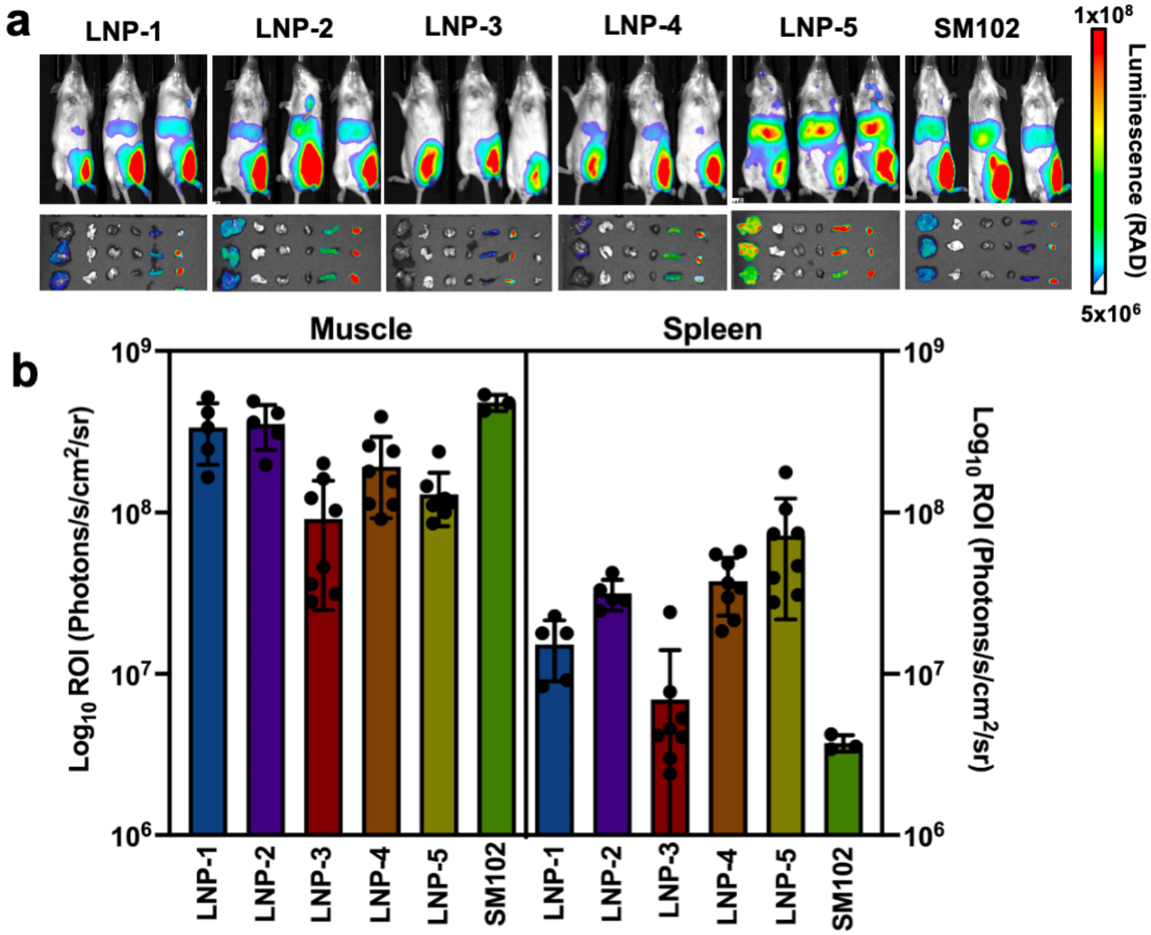


***Figure S1*. I.m. administration of Top LNPs. a)** representative images of bioluminescence in vivo and ex vivo 6 post LNPs administration (2 µg Fluc circRNA per mouse). **b)** quantification of bioluminescent signal from muscles and spleens. Data are presented as Mean±SD (n=5 for LNP-1 and LNP-2, n=8 for LNP-3, LNP-4 and LNP-5, n=3 for SM102).

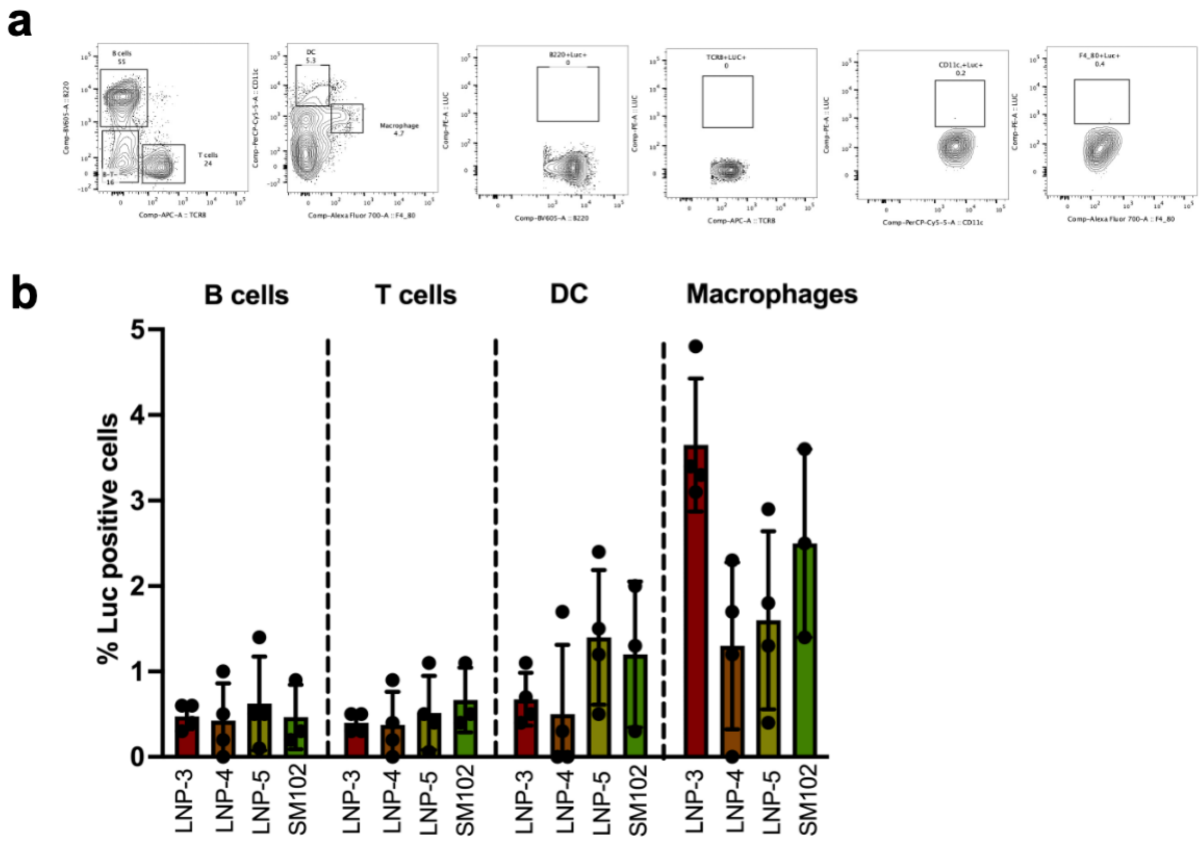


***Figure S2*. Flow cytometry analysis of mice spleens 6 hr post administration of LNPs with 4 µg Fluc circRNA per mouse.** a) workflow of flow cytometry gating strategy for spleen tissue. b) quantification of Luc+ cells in the spleen in different cell types. Data are presented as Means ± SD (n = 4 biological replicates for LNP-3, LNP-4 and LNP-5 and n=3 for SM102).

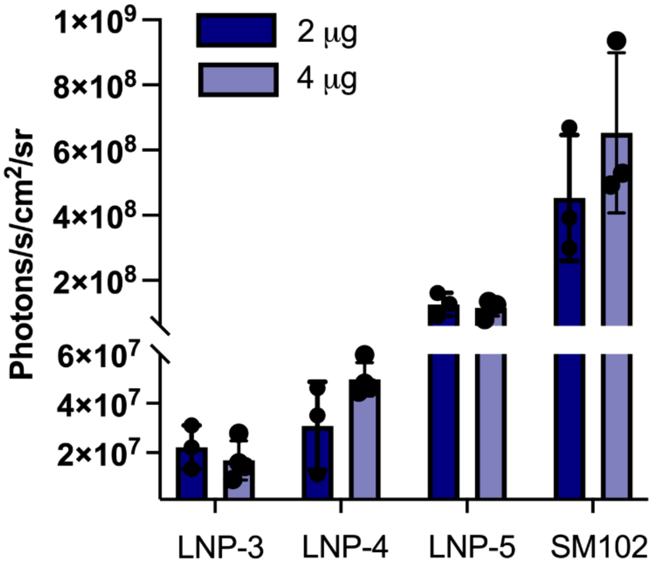


***Figure S3***. Dose dependence of ex vivo bioluminescent signal in liver (BALB/c mice) 6 hr post r.o. administration of Fluc circRNA. Data are presented as Means ± SD (n = 4 biological replicates for LNP-3, LNP-4 and LNP-5 and n=3 for SM102).

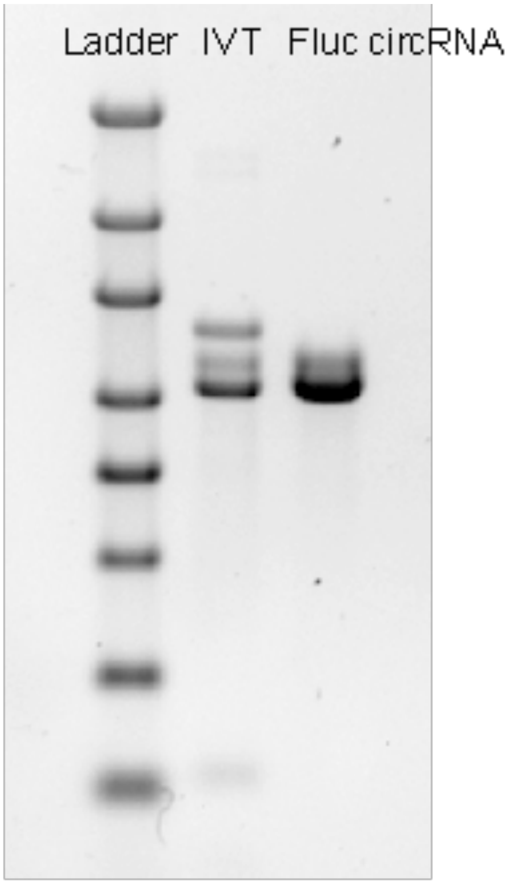


***Figure S4.*** Denaturing 0.8% agarose gel electrophoresis in 1x NorthernMax buffer of IVT product and Fluc circRNA after HPLC purification. See Supporting Information Table S3 for the sequence of Fluc circRNA.

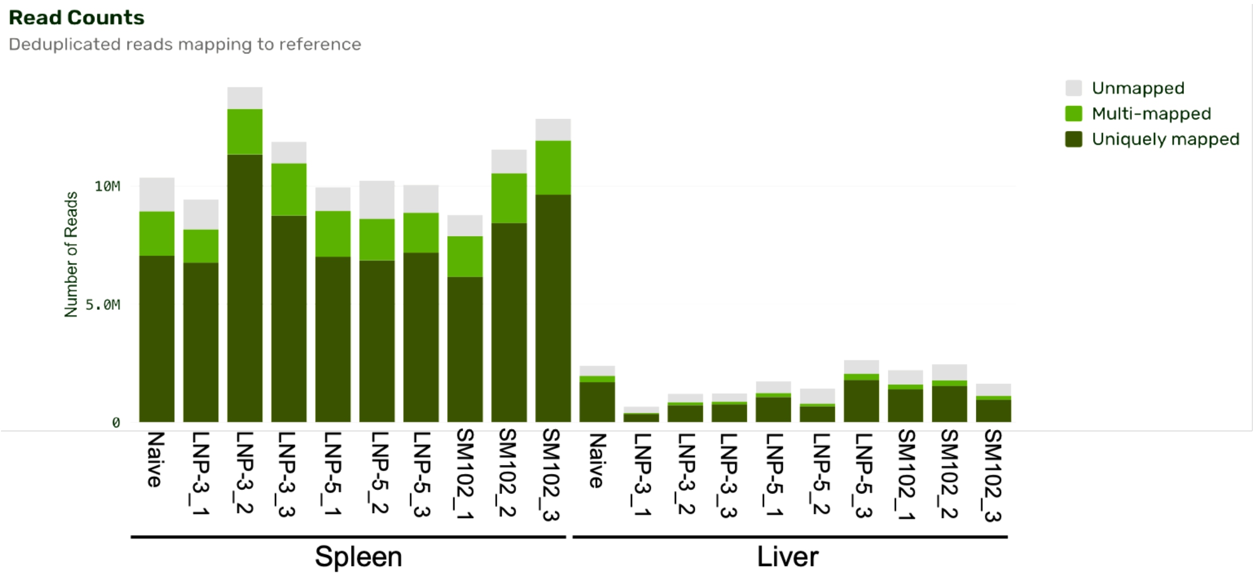


***Figure S5.*** Summary of sequencing read alignment and deduplication metrics.

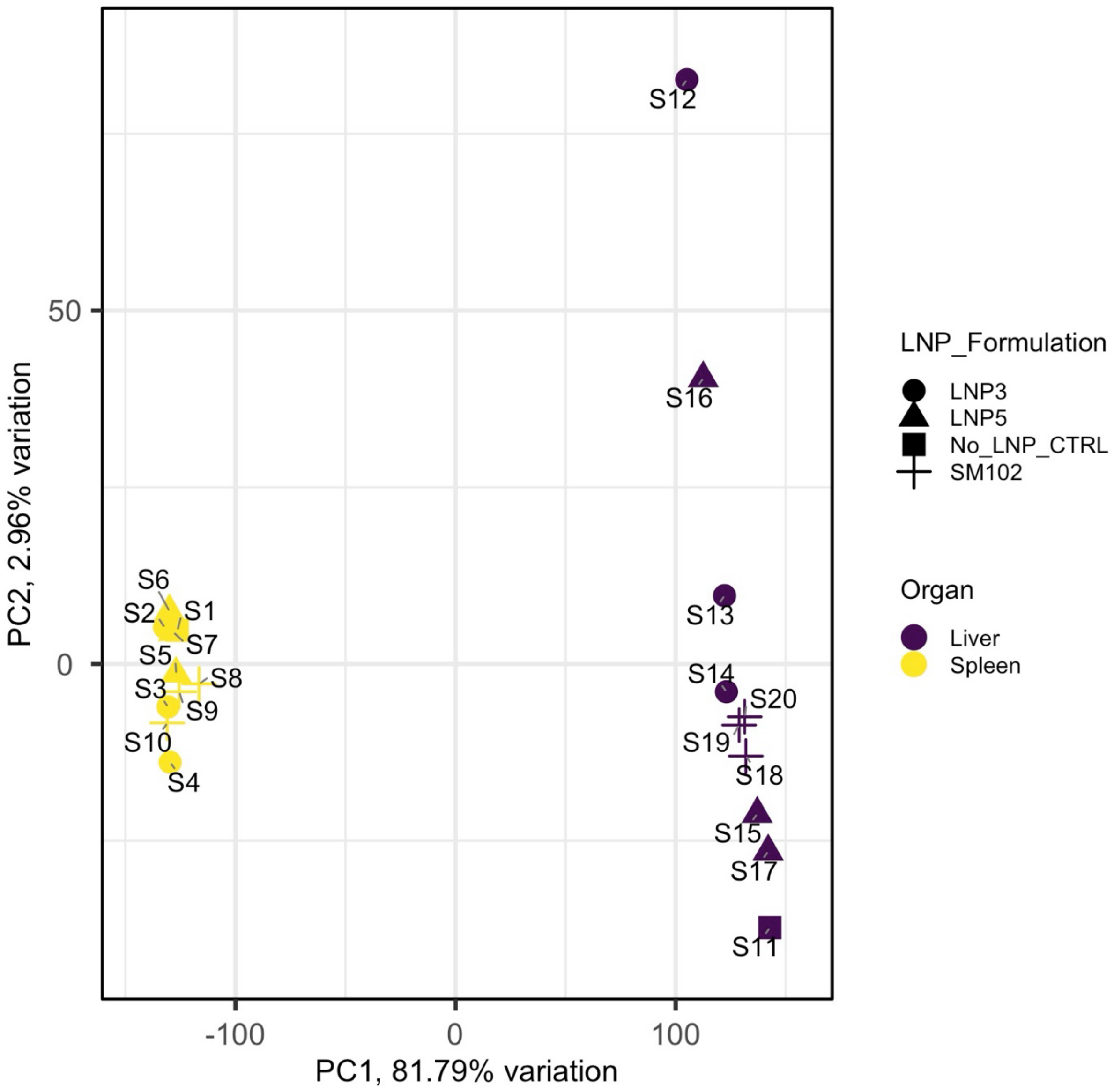


***Figure S6.*** Principle Component Analysis. Data capture organ specific variability wrt. transcript expression.

***Table S1***. Critical quality attributes (CQAs) of LNP screening library and their corresponding functional readouts in T cells and Jurkat cells. (a) Plate map layout of DOE designed LNPs (1–54) and Cytiva control LNPs across Plates 1–4. (b) Hydrodynamic size (nm) for all LNPs. (c) Polydispersity index (PDI) values (d) Encapsulation efficiency (%EE) of the circRNA in LNPs. (e) Jurkat cell screening results, presented as luminescence/$EC_{50}$. (f) Primary T cell screening, expressed as luminescence/$EC_{50}$.

| (a) Plate map | | | | | | | | | |
|---|---|---|---|---|---|---|---|---|---|
| Plate-1 | C1 | 7 | 15 | 23 | 31 | 39 | 47 | C11 | C8 |
| | C5 | 8 | 16 | 24 | 32 | 40 | 48 | C12 | C9 |
| | 1 | 9 | 17 | 25 | 33 | 41 | 49 | C2 | C10 |
| | 2 | 10 | 18 | 26 | 34 | 42 | 50 | C3 | C1-2 |
| | 3 | 11 | 19 | 27 | 35 | 43 | 51 | C4 | C5-2 |
| | 4 | 12 | 20 | 28 | 36 | 44 | 52 | C5 | |
| | 5 | 13 | 21 | 29 | 37 | 45 | 53 | C6 | |
| | 6 | 14 | 22 | 30 | 38 | 46 | 54 | C7 | |
| Plate-2 | C1 | 7 | 15 | 23 | 31 | 39 | 47 | C2 | C1-2 |
| | C5 | 8 | 16 | 24 | 32 | 40 | 48 | C3 | C5-2 |
| | 1 | 9 | 17 | 25 | 33 | 41 | 49 | C4 | |
| | 2 | 10 | 18 | 26 | 34 | 42 | 50 | C6 | |
| | 3 | 11 | 19 | 27 | 35 | 43 | 51 | C7 | |
| | 4 | 12 | 20 | 28 | 36 | 44 | 52 | C8 | |
| | 5 | 13 | 21 | 29 | 37 | 45 | 53 | C9 | |
| | 6 | 14 | 22 | 30 | 38 | 46 | 54 | C10 | |
| Plate-3 | C1 | 7 | 15 | 23 | 31 | 39 | 47 | C2 | C10 |
| | C5 | 8 | 16 | 24 | 32 | 40 | 48 | C3 | C1-2 |
| | 1 | 9 | 17 | 25 | 33 | 41 | 49 | C4 | C5-2 |
| | 2 | 10 | 18 | 26 | 34 | 42 | 50 | C5 | |
| | 3 | 11 | 19 | 27 | 35 | 43 | 51 | C6 | |
| | 4 | 12 | 20 | 28 | 36 | 44 | 52 | C7 | |
| | 5 | 13 | 21 | 29 | 37 | 45 | 53 | C8 | |
| | 6 | 14 | 22 | 30 | 38 | 46 | 54 | C9 | |
| Plate-4 | C1 | 7 | 15 | 23 | 31 | 39 | 47 | C2 | C1-2 |
| | C5 | 8 | 16 | 24 | 32 | 40 | 48 | C3 | C5-2 |
| | 1 | 9 | 17 | 25 | 33 | 41 | 49 | C4 | |
| | 2 | 10 | 18 | 26 | 34 | 42 | 50 | C6 | |
| | 3 | 11 | 19 | 27 | 35 | 43 | 51 | C7 | |
| | 4 | 12 | 20 | 28 | 36 | 44 | 52 | C8 | |
| | 5 | 13 | 21 | 29 | 37 | 45 | 53 | C9 | |
| | 6 | 14 | 22 | 30 | 38 | 46 | 54 | C10 | |

| | (b) Size (nm) | | | | | | | | |
|---|---|---|---|---|---|---|---|---|---|
| Plate-1 | 74.7 | 62.8 | 75.3 | 65.6 | 97.2 | 79.4 | 91.9 | 117.1 | 102.2 |
| | 102.5 | 61.6 | 54.7 | 130.7 | 59.4 | 76.2 | 73.8 | 81.5 | 70.5 |
| | 81.6 | 61.6 | 59.3 | 53.0 | 99.1 | 60.4 | 97.3 | 115.7 | 77.3 |
| | 79.6 | 60.1 | 59.2 | 69.5 | 54.9 | 109.9 | 64.8 | 114.3 | 69.7 |
| | 72.5 | 51.8 | 81.0 | 58.5 | 58.1 | 70.0 | 91.2 | 87.7 | 91.8 |
| | 71.4 | 70.3 | 114.6 | 60.0 | 63.6 | 64.2 | 64.2 | 94.1 | |
| | 71.7 | 70.6 | 62.0 | 57.3 | 104.6 | 83.1 | 87.0 | 66.5 | |
| | 87.1 | 62.0 | 74.0 | 81.6 | 105.8 | 82.1 | 79.0 | 107.3 | |
| Plate-2 | 74.9 | 51.5 | 79.1 | 57.0 | 67.9 | 63.6 | 75.9 | 157.0 | 70.6 |
| | 94.9 | 67.5 | 59.7 | 96.8 | 59.3 | 69.8 | 88.3 | 101.6 | 87.5 |
| | 93.0 | 64.5 | 65.8 | 57.3 | 106.1 | 68.0 | 100.9 | 83.0 | |
| | 113.5 | 68.8 | 62.6 | 62.5 | 64.9 | 131.1 | 56.9 | 68.8 | |
| | 83.1 | 74.4 | 78.1 | 56.7 | 63.1 | 58.0 | 94.7 | 93.9 | |
| | 77.8 | 85.7 | 101.3 | 57.5 | 61.6 | 64.2 | 57.6 | 103.4 | |
| | 71.3 | 87.2 | 62.6 | 66.7 | 78.0 | 60.7 | 60.8 | 72.6 | |
| | 127.1 | 57.5 | 78.2 | 81.6 | 121.8 | 61.4 | 59.5 | 67.8 | |
| Plate-3 | 66.7 | 71.9 | 109.6 | 75.2 | 99.7 | 106.0 | 113.2 | 134.5 | 75.0 |
| | 93.1 | 74.5 | 61.7 | 133.4 | 82.1 | 83.3 | 93.4 | 125.4 | 68.3 |
| | 99.5 | 76.3 | 72.1 | 78.6 | 158.3 | 74.4 | 118.0 | 67.1 | 118.1 |
| | 92.5 | 72.4 | 73.6 | 84.8 | 77.8 | 125.7 | 75.7 | 92.3 | |
| | 100.1 | 73.0 | 84.0 | 81.8 | 89.2 | 74.3 | 116.6 | 67.5 | |
| | 80.6 | 87.7 | 132.7 | 81.6 | 79.2 | 71.4 | 69.0 | 94.1 | |
| | 85.3 | 99.4 | 98.9 | 185.0 | 87.8 | 61.5 | 67.4 | 100.6 | |
| | 117.3 | 65.7 | 100.5 | 119.4 | 107.2 | 74.4 | 69.8 | 90.0 | |
| Plate-4 | 76.2 | 53.1 | 93.3 | 123.8 | 107.8 | 70.1 | 72.1 | 119.0 | 75.2 |
| | 90.4 | 60.0 | 56.7 | 85.6 | 62.3 | 85.9 | 104.7 | 119.7 | 87.8 |
| | 67.8 | 54.3 | 53.8 | 66.8 | 65.9 | 94.3 | 122.2 | 77.2 | |
| | 76.6 | 53.2 | 53.8 | 94.0 | 70.1 | 124.6 | 64.1 | 74.8 | |
| | 60.8 | 47.1 | 91.3 | 64.2 | 64.6 | 57.7 | 91.5 | 95.1 | |
| | 74.1 | 73.6 | 84.7 | 66.6 | 67.2 | 72.8 | 63.9 | 104.4 | |
| | 53.4 | 57.5 | 75.9 | 79.2 | 94.1 | 67.4 | 72.5 | 72.4 | |
| | 75.1 | 56.1 | 64.5 | 78.5 | 136.9 | 73.0 | 66.6 | 72.0 | |

| (c) PDI | | | | | | | | | |
|---|---|---|---|---|---|---|---|---|---|
| Plate-1 | 0.32 | 1.00 | 0.22 | 0.23 | 0.15 | 0.39 | 1.00 | 0.09 | 0.23 |
| | 0.33 | 0.25 | 0.34 | 0.17 | 0.29 | 0.32 | 0.18 | 0.31 | 0.31 |
| | 0.17 | 0.24 | 0.19 | 0.37 | 0.11 | 0.44 | 0.09 | 0.27 | 0.29 |
| | 0.18 | 0.30 | 0.22 | 0.19 | 0.29 | 0.08 | 0.46 | 0.43 | 0.23 |
| | 0.35 | 0.12 | 0.25 | 0.22 | 0.37 | 0.55 | 0.17 | 0.16 | 0.23 |
| | 0.38 | 0.25 | 0.15 | 0.24 | 0.32 | 0.32 | 0.47 | 0.29 | |
| | 0.40 | 0.13 | 0.20 | 0.39 | 0.44 | 0.48 | 0.47 | 0.16 | |
| | 0.22 | 0.41 | 0.19 | 0.09 | 0.28 | 0.51 | 0.45 | 0.31 | |
| Plate-2 | 0.35 | 0.46 | 0.08 | 0.31 | 0.17 | 0.21 | 0.13 | 0.41 | 0.20 |
| | 0.36 | 0.23 | 0.41 | 0.16 | 0.28 | 0.16 | 0.11 | 0.44 | 0.26 |
| | 0.24 | 0.29 | 0.33 | 1.00 | 0.07 | 0.25 | 0.15 | 0.31 | |
| | 0.30 | 0.30 | 0.35 | 0.18 | 0.44 | 0.15 | 0.27 | 0.18 | |
| | 1.00 | 0.41 | 0.13 | 0.27 | 0.31 | 0.49 | 0.06 | 0.25 | |
| | 0.35 | 0.12 | 0.13 | 0.27 | 0.23 | 0.20 | 0.21 | 0.19 | |
| | 0.38 | 0.16 | 0.21 | 0.30 | 0.25 | 0.32 | 0.39 | 0.38 | |
| | 0.13 | 0.26 | 0.13 | 0.20 | 0.24 | 0.43 | 0.23 | 0.22 | |
| Plate-3 | 0.45 | 0.31 | 0.19 | 0.36 | 0.49 | 0.52 | 1.00 | 0.41 | 0.25 |
| | 0.37 | 0.30 | 0.39 | 0.22 | 0.30 | 0.19 | 0.20 | 0.39 | 0.46 |
| | 0.20 | 0.43 | 0.36 | 1.00 | 0.19 | 1.00 | 0.42 | n/a | 0.22 |
| | 0.45 | 0.38 | 0.40 | 0.39 | 0.35 | 0.22 | 0.43 | n/a | |
| | 0.39 | 0.50 | 0.32 | 0.37 | 0.35 | 1.00 | 0.27 | 0.44 | |
| | 0.36 | 0.31 | 0.26 | 0.39 | 0.23 | 0.32 | n/a | 0.21 | |
| | 1.00 | 0.31 | 0.40 | 1.00 | 0.42 | 0.32 | 0.43 | 0.37 | |
| | 0.17 | 0.27 | 0.11 | 0.18 | 0.16 | 0.30 | 0.24 | * | |
| Plate-4 | 0.14 | 0.55 | 0.04 | 0.28 | 1.00 | 0.17 | 0.23 | 0.28 | 0.21 |
| | 0.26 | 0.25 | 0.31 | 0.37 | 0.27 | 0.10 | 0.23 | 0.39 | 0.17 |
| | 0.17 | 0.33 | 0.29 | 0.25 | 0.17 | 0.23 | 0.48 | n/a | |
| | 0.14 | 0.21 | 0.22 | 0.19 | 0.39 | 0.30 | 0.20 | 0.21 | |
| | 0.19 | 0.27 | 0.23 | 0.25 | 0.27 | 0.21 | 0.07 | 0.30 | |
| | 0.09 | 0.23 | 0.31 | 0.31 | 0.40 | 0.16 | 0.20 | 0.22 | |
| | 0.21 | 0.11 | 0.38 | 0.22 | 0.20 | 0.18 | 0.28 | 0.45 | |
| | 0.17 | 0.15 | 0.23 | 0.23 | 0.49 | 0.19 | 0.16 | 0.27 | |

** Multimodal notification, value not available*

| | (d) %EE | | | | | | | | |
|---|---|---|---|---|---|---|---|---|---|
| Plate-1 | 98.0 | 98.6 | 98.3 | 95.6 | 67.5 | 95.7 | 92.5 | 97.9 | 85.7 |
| | 85.6 | 98.3 | 97.8 | 60.8 | 98.0 | 97.8 | 97.5 | 97.4 | 97.6 |
| | 98.1 | 98.4 | 87.3 | 97.8 | 99.3 | 97.4 | 79.5 | 92.3 | 96.9 |
| | 98.9 | 98.4 | 97.7 | 97.9 | 97.5 | 77.0 | 97.2 | 89.5 | 97.8 |
| | 97.5 | 98.6 | 95.2 | 97.5 | -10.1 | 96.2 | 97.0 | 98.5 | 87.2 |
| | 97.4 | 97.8 | 90.6 | 94.3 | 97.9 | 98.8 | 97.9 | 88.5 | |
| | 88.7 | 81.5 | 97.8 | 97.7 | 92.2 | 96.8 | 97.2 | 96.2 | |
| | 81.8 | 97.7 | 98.5 | 97.2 | 96.6 | 96.8 | 97.2 | 80.0 | |
| Plate-2 | 96.7 | 97.8 | 99.3 | 95.8 | 92.6 | 98.7 | 98.2 | 89.8 | 97.7 |
| | 88.4 | 98.2 | 98.0 | 84.6 | 98.4 | 98.6 | 97.5 | 92.0 | 92.0 |
| | 93.7 | 97.8 | 98.0 | 98.0 | 99.5 | 96.6 | 74.0 | 98.6 | |
| | 95.3 | 97.9 | 98.2 | 98.4 | 98.6 | 63.5 | 98.6 | 96.7 | |
| | 96.5 | 97.8 | 96.2 | 98.2 | 98.3 | 97.8 | 99.5 | 86.9 | |
| | 98.2 | 97.8 | 96.8 | 98.1 | 98.2 | 98.8 | 98.6 | 83.9 | |
| | 95.2 | 86.4 | 98.0 | 97.3 | 96.7 | 98.3 | 98.5 | 97.8 | |
| | 71.0 | 98.2 | 98.5 | 98.4 | 90.9 | 98.3 | 98.4 | 98.6 | |
| Plate-3 | 98.4 | 97.5 | 98.0 | 89.2 | 76.0 | 98.7 | 98.3 | 93.3 | 98.3 |
| | 91.5 | 97.6 | 98.4 | 55.1 | 97.8 | 98.4 | 97.4 | 91.5 | 98.1 |
| | 95.7 | 96.5 | 97.6 | 98.4 | 67.4 | 98.4 | 73.7 | 95.8 | 88.2 |
| | 95.9 | 96.4 | 97.7 | 97.8 | 97.4 | 64.3 | 98.3 | 90.5 | |
| | 98.7 | 97.9 | 93.1 | 97.2 | 96.6 | 98.4 | 98.7 | 98.1 | |
| | 97.9 | 97.0 | 83.0 | 96.9 | 97.2 | 98.7 | 98.3 | 82.2 | |
| | 97.9 | 82.5 | 97.6 | 97.5 | 97.2 | 98.5 | 98.4 | 80.3 | |
| | 74.9 | 98.6 | 98.0 | 93.1 | 93.0 | 98.5 | 98.3 | 97.1 | |
| Plate-4 | 97.6 | 98.2 | 99.7 | 96.6 | 93.2 | 98.3 | 92.7 | 93.4 | 96.9 |
| | 91.6 | 98.3 | 98.4 | 94.7 | 98.1 | 97.5 | 91.9 | 90.8 | 91.1 |
| | 96.6 | 97.9 | 98.5 | 98.1 | 97.2 | 90.4 | 79.5 | 95.2 | |
| | 98.4 | 98.0 | 98.5 | 88.7 | 97.3 | 74.6 | 98.0 | 97.3 | |
| | 97.6 | 96.7 | 98.0 | 97.6 | 98.0 | 97.8 | 99.5 | 85.1 | |
| | 98.4 | 98.1 | 97.1 | 97.2 | 97.8 | 97.5 | 96.5 | 79.6 | |
| | 97.7 | 95.0 | 93.9 | 98.4 | 93.9 | 97.1 | 97.2 | 97.9 | |
| | 92.9 | 98.8 | 98.1 | 96.0 | 87.3 | 97.3 | 97.2 | 98.6 | |

| | (e) Screening in Jurkat cells Luminescence/EC50 | | | | | | | | |
|---|---|---|---|---|---|---|---|---|---|
| Plate-1 | 189.9 | 37.5 | 392.4 | 99.9 | 65.8 | 165.1 | 61.2 | 971 | 629.2 |
| | 482.7 | 226.7 | 118.7 | 144.7 | 283.8 | 137.5 | 127.9 | 423.6 | 167.1 |
| | 139.4 | 261.9 | 244.4 | 60.1 | 322.4 | 198.1 | 42.2 | 44.3 | 802.8 |
| | 323.6 | 247.7 | 239.4 | 573.4 | 115.5 | 27.4 | 117.6 | 66.9 | 220.5 |
| | 161.5 | 50.5 | 347.7 | 506.5 | 36.4 | 73.7 | 120.4 | 262.4 | 446.7 |
| | 421.5 | 458.7 | 828.3 | 506.1 | 452.4 | 140.8 | 83.3 | 565.7 | |
| | 48.3 | 59 | 485.1 | 82.4 | 117.6 | 304 | 329.5 | 60.3 | |
| | 80.3 | 223.5 | 899.8 | 1343.6 | 121.9 | 309.4 | 308 | 907.9 | |
| Plate-2 | 305 | 146.6 | 1420.5 | 85.5 | 207.4 | 641.1 | 89.3 | 20.4 | 455.2 |
| | 825.4 | 1296 | 444.1 | 796.8 | 341.1 | 525.3 | 1241.8 | 29.9 | 1015 |
| | 267.2 | 817.8 | 963.8 | 109.1 | 187.4 | 114.3 | 89.4 | 190.8 | |
| | 542 | 810.1 | 911.1 | 396.6 | 141.9 | 203.3 | 526.4 | 86.3 | |
| | 876.8 | 126.9 | 376 | 219 | 220.4 | 105 | 140.7 | 1168.4 | |
| | 892.3 | 1637.4 | 1344.5 | 326.1 | 247.4 | 1066.2 | 571.6 | 1087.2 | |
| | 96.9 | 774.9 | 568.7 | 92.4 | 334.5 | 809.3 | 848.7 | 123.9 | |
| | 292 | 691.7 | 657.6 | 1285.7 | 704.3 | 818.4 | 880.4 | 615 | |
| Plate-3 | 211.8 | 198.1 | 433.9 | 154.8 | 160.9 | 882.8 | 82.6 | 35.6 | 754.7 |
| | 873.6 | 574.4 | 166 | 133.1 | 145.7 | 728.7 | 707.1 | 47.1 | 217.1 |
| | 544.2 | 842.9 | 614.8 | 87 | 37.3 | 81.7 | 409.4 | 129 | 716.8 |
| | 118.9 | 820.6 | 896.8 | 300.1 | 265.9 | 188.7 | 782.1 | 718.9 | |
| | 493.9 | 112.3 | 493.4 | 556.7 | 738.8 | 152 | 817.8 | 257 | |
| | 500.4 | 199.8 | 632.7 | 631.5 | 774.7 | 889.2 | 279 | 1058.3 | |
| | 54.4 | 408.3 | 469.1 | 101.5 | 244.3 | 810.6 | 680.1 | 1084.2 | |
| | 178.6 | 365.3 | 591.9 | 1037.8 | 186.9 | 648.7 | 608.4 | 104.6 | |
| Plate-4 | 297.6 | 63.6 | 469.9 | 112.8 | 129.5 | 326.6 | 96.9 | 28 | 303.4 |
| | 879.9 | 539 | 202.5 | 644.3 | 604 | 161.1 | 1232.5 | 60.9 | 692.1 |
| | 166.8 | 307.7 | 310.7 | 422.5 | 662.5 | 351.5 | 236.4 | 147.2 | |
| | 1261 | 266.9 | 257 | 520.3 | 435.2 | 452.4 | 642.9 | 310.4 | |
| | 332.2 | 81.2 | 208.9 | 491 | 542 | 743 | 559.4 | 1336.4 | |
| | 1325.8 | 1476 | 841 | 539.3 | 584.1 | 897.1 | 1186 | 795.5 | |
| | 113 | 246.7 | 326 | 94.7 | 481 | 1547 | 2167.7 | 181.2 | |
| | 894.9 | 441.5 | 711.8 | 996.3 | 2368.8 | 2452.3 | 2515.1 | 759.4 | |

| | (f) Screening in T cells Luminescence/EC50 | | | | | | | | |
|---|---|---|---|---|---|---|---|---|---|
| Plate-1 | 4.1 | 0.5 | 6.6 | 2.1 | 0.6 | 1 | 0.6 | 10.7 | 11.6 |
| | 4 | 3.2 | 1 | 1.2 | 5.5 | 0.6 | 0.8 | 6 | 4.2 |
| | 1.4 | 3.9 | 3.8 | 0.9 | 1.9 | 0.5 | 0.2 | 0.7 | 12.7 |
| | 3.7 | 3.5 | 3.6 | 5 | 0.9 | 0.2 | 0.5 | 1 | 4.5 |
| | 2.2 | 0.6 | 9 | 5.8 | 0.4 | 0.4 | 0.7 | 2.4 | 2.3 |
| | 6.8 | 3.9 | 4.3 | 5.3 | 5.4 | 0.8 | 0.3 | 2.2 | |
| | 1.3 | 0.7 | 5.4 | 0.7 | 0.5 | 1 | 0.9 | 0.5 | |
| | 0.7 | 4.7 | 8.5 | 6.5 | 0.5 | 1.3 | 1.1 | 2.8 | |
| Plate-2 | 3.4 | 0.8 | 11 | 0.8 | 1 | 16.9 | 0.7 | 0.3 | 3.9 |
| | 2.2 | 5 | 2.4 | 2.2 | 1.6 | 7.2 | 7 | 0.7 | 1.2 |
| | 8.1 | 7.7 | 8.4 | 0.5 | 1.4 | 2.2 | 0.7 | 2.1 | |
| | 3.4 | 9.3 | 8.1 | 1.3 | 0.4 | 1 | 4 | 0.4 | |
| | 17.9 | 1.3 | 7.1 | 1.3 | 1.1 | 0.4 | 3.2 | 1.5 | |
| | 9.2 | 7.1 | 8.5 | 1.5 | 1.7 | 1.9 | 1.6 | 2.4 | |
| | 1.4 | 1.2 | 10.2 | 0.9 | 8.4 | 5.8 | 6 | 3.4 | |
| | 2.4 | 5.5 | 5.2 | 8.1 | 5.2 | 7.9 | 4.8 | 3.3 | |
| Plate-3 | 3 | 2.2 | 3.5 | 2 | 0.8 | 15.6 | 0.8 | 0.4 | 8.3 |
| | 2.4 | 2.4 | 1.2 | 0.5 | 2.6 | 10.7 | 10.1 | 0.8 | 2.9 |
| | 12.5 | 4.6 | 3.9 | 0.8 | 0.4 | 1.1 | 1.8 | 0.4 | 1.3 |
| | 1.6 | 4.2 | 4.1 | 3.8 | 1.2 | 3.1 | 11.6 | 1.3 | |
| | 7.8 | 0.7 | 2.7 | 4.5 | 5.1 | 1.6 | 8.5 | 3.2 | |
| | 4.9 | 2.1 | 1.2 | 4.9 | 4.2 | 7.7 | 2 | 1.4 | |
| | 0.6 | 1.1 | 7.9 | 0.8 | 5.2 | 10 | 9.6 | 3.2 | |
| | 2.1 | 8 | 4.6 | 6.6 | 3.7 | 11 | 10.5 | 3.1 | |
| Plate-4 | 3.3 | 0.3 | 1.9 | 1.2 | 0.5 | 6.7 | 0.9 | 0.3 | 4.1 |
| | 2.4 | 1.4 | 0.6 | 1.8 | 7.6 | 3.8 | 7.3 | 1 | 1.3 |
| | 3.3 | 2.2 | 2.3 | 5.7 | 4.4 | 4.9 | 0.7 | 0.4 | |
| | 8.4 | 2.3 | 2.1 | 3.7 | 3.9 | 1.3 | 7.6 | 3.4 | |
| | 4.4 | 0.6 | 1.5 | 5.1 | 5.9 | 8.3 | 4 | 1.3 | |
| | 11 | 6.2 | 3.4 | 5.2 | 5.2 | 4.7 | 6.9 | 1.7 | |
| | 1.7 | 0.9 | 6 | 1.2 | 4.6 | 12.3 | 12.3 | 3.4 | |
| | 1.7 | 1 | 5.8 | 8.1 | 6.9 | 14 | 12.1 | 5 | |

***Table S2***. Cytokines screening in serum from BALB/c mice 6 hr post r.o. administration of LNP-3, LNP-4, LNP-5 and SM102. Fluc circRNA dose was 2 µg per mouse. n=3 biological replicates. Data are shown as concentration pg/ml.

| | Naïve 1 | Naïve 2 | Naïve 3 | LNP-3 1 | LNP-3 2 | LNP-3 3 | LNP-4 1 | LNP-4 2 | LNP-4 3 | LNP-5 1 | LNP-5 2 | LNP-5 3 | SM102 1 | SM102 2 | SM102 3 |
|---|---|---|---|---|---|---|---|---|---|---|---|---|---|---|---|
| BAFF/TNFSF13B | 119.8 | 113.9 | 106.2 | 119.2 | 111.2 | 156.6 | 123.1 | 112.4 | 112.1 | 112.0 | 109.4 | 110.8 | 103.4 | 129.8 | 106.0 |
| BTC | 3.2 | 4.2 | 1.0 | 6.1 | 4.7 | 3.9 | 1.0 | 4.6 | 4.6 | 3.5 | 3.4 | 1.0 | 4.4 | 4.1 | 5.3 |
| ENA-78/CXCL5 | 46.3 | 72.5 | 46.0 | 47.4 | 61.3 | 35.7 | 55.0 | 49.6 | 61.0 | 55.8 | 66.7 | 86.8 | 61.7 | 45.5 | 55.3 |
| EOTAXIN/CCL11 | 483.5 | 577.4 | 481.4 | 778.1 | 873.2 | 666.8 | 696.3 | 580.4 | 719.0 | 566.8 | 510.7 | 427.2 | 504.1 | 541.6 | 467.7 |
| G-CSF/CSF-3 | 3.2 | 7.2 | 1.0 | 56.3 | 41.4 | 29.6 | 12.0 | 7.7 | 13.1 | 9.8 | 5.8 | 3.5 | 6.7 | 1.0 | 7.2 |
| GM-CSF | 1.0 | 1.0 | 1.0 | 3.7 | 1.0 | 3.4 | 1.0 | 1.0 | 1.0 | 1.0 | 1.0 | 1.0 | 3.4 | 1.0 | 1.0 |
| GROa/KC/CXCL1 | 1.0 | 17.3 | 1.0 | 48.4 | 101.0 | 37.9 | 16.3 | 10.6 | 1.0 | 23.2 | 1.0 | 5.6 | 1.0 | 1.0 | 1.0 |
| IFNa | 2.3 | 2.6 | 2.4 | 3.1 | 2.9 | 3.2 | 2.7 | 2.7 | 2.5 | 3.1 | 2.9 | 2.4 | 3.2 | 2.8 | 4.5 |
| IFNg | 1.0 | 1.0 | 1.0 | 1.0 | 1.0 | 1.0 | 1.0 | 1.0 | 1.0 | 1.0 | 1.0 | 1.0 | 1.0 | 1.0 | 1.0 |
| IL-10 | 1.0 | 1.0 | 1.0 | 1.0 | 1.0 | 1.0 | 1.0 | 1.0 | 1.0 | 1.0 | 1.0 | 1.0 | 1.0 | 1.0 | 1.0 |
| IL-12P70 | 3.6 | 3.5 | 3.0 | 4.8 | 3.8 | 4.4 | 3.7 | 3.6 | 3.9 | 4.1 | 4.0 | 3.8 | 4.1 | 4.1 | 4.5 |
| IL-13 | 4.5 | 1.0 | 1.0 | 1.0 | 1.0 | 3.7 | 1.0 | 1.0 | 1.0 | 1.0 | 1.0 | 1.0 | 1.0 | 1.0 | 1.0 |
| IL-15/IL-15R | 1.0 | 1.0 | 1.0 | 1.0 | 1.0 | 1.0 | 1.0 | 1.0 | 1.0 | 1.0 | 1.0 | 1.0 | 1.0 | 1.0 | 1.0 |
| IL-17A/CTLA8 | 1.0 | 1.0 | 1.0 | 6.1 | 5.6 | 7.7 | 4.7 | 5.0 | 5.1 | 5.4 | 4.5 | 1.0 | 6.3 | 5.2 | 5.2 |
| IL-18 | 5.2 | 10.4 | 4.4 | 17.0 | 12.6 | 15.5 | 11.1 | 7.6 | 9.8 | 11.9 | 10.2 | 9.9 | 9.8 | 8.9 | 10.3 |
| IL-19 | 35.7 | 31.1 | 1.0 | 34.3 | 34.7 | 52.9 | 34.5 | 17.9 | 51.9 | 41.6 | 47.7 | 18.7 | 41.9 | 32.4 | 48.5 |
| IL-1a | 2.7 | 3.8 | 2.6 | 6.6 | 5.0 | 6.1 | 4.3 | 4.4 | 4.3 | 6.7 | 4.3 | 5.7 | 4.5 | 4.2 | 4.9 |
| IL-1b | 3.8 | 5.3 | 1.0 | 5.3 | 4.7 | 6.8 | 4.3 | 4.1 | 5.2 | 5.9 | 5.9 | 4.0 | 6.0 | 4.6 | 5.0 |
| IL-2 | 1.0 | 2.8 | 8.6 | 4.5 | 3.8 | 3.5 | 1.0 | 4.7 | 3.3 | 5.0 | 4.7 | 3.1 | 3.3 | 3.1 | 3.0 |
| IL-22 | 1.0 | 1.0 | 1.0 | 1.0 | 1.0 | 1.0 | 1.0 | 1.0 | 1.0 | 1.0 | 1.0 | 1.0 | 1.0 | 1.0 | 1.0 |
| IL-23 | 1.0 | 8.6 | 1.0 | 11.2 | 8.2 | 17.0 | 10.1 | 1.0 | 1.0 | 10.3 | 18.6 | 1.0 | 15.4 | 10.0 | 12.0 |
| IL-25/IL-17 | 1.0 | 1.0 | 1.0 | 1.0 | 1.0 | 1.0 | 1.0 | 1.0 | 1.0 | 1.0 | 1.0 | 1.0 | 1.0 | 1.0 | 1.0 |
| IL-27 | 13.9 | 4.4 | 8.1 | 1.0 | 1.0 | 24.3 | 1.0 | 8.1 | 6.0 | 18.7 | 3.6 | 6.2 | 5.4 | 6.8 | 1.0 |
| IL-28 | 1.0 | 1.0 | 1.0 | 1.0 | 1.0 | 1.0 | 1.0 | 1.0 | 1.0 | 1.0 | 1.0 | 1.0 | 1.0 | 1.0 | 1.0 |
| IL-2RA | 19.1 | 15.7 | 20.7 | 18.2 | 16.4 | 40.1 | 27.5 | 24.6 | 17.4 | 18.7 | 22.8 | 24.4 | 24.1 | 15.3 | 20.6 |
| IL-3 | 1.0 | 1.0 | 1.0 | 1.0 | 1.0 | 1.0 | 1.0 | 1.0 | 1.0 | 1.0 | 1.0 | 1.0 | 1.0 | 1.0 | 1.0 |
| IL-31 | 1.0 | 1.0 | 1.0 | 1.0 | 1.0 | 1.0 | 1.0 | 1.0 | 1.0 | 1.0 | 1.0 | 1.0 | 1.0 | 1.0 | 1.0 |
| IL-33 | 1.0 | 1.0 | 1.0 | 0.9 | 1.0 | 0.8 | 1.0 | 1.0 | 1.0 | 1.0 | 1.0 | 1.0 | 1.0 | 1.0 | 1.0 |
| IL-4 | 1.0 | 1.0 | 1.0 | 1.0 | 3.3 | 3.7 | 3.1 | 1.0 | 1.0 | 1.0 | 1.0 | 4.2 | 1.0 | 1.0 | 1.0 |
| IL-5 | 1.0 | 1.0 | 1.0 | 46.2 | 114.7 | 44.6 | 36.7 | 4.8 | 25.5 | 51.0 | 3.8 | 5.0 | 1.0 | 1.0 | 1.0 |
| IL-6 | 1.0 | 1.0 | 1.0 | 49.8 | 80.1 | 34.9 | 8.0 | 4.1 | 6.2 | 14.4 | 1.0 | 1.0 | 1.0 | 1.0 | 1.0 |
| IL-7 | 24.5 | 7.6 | 12.6 | 12.6 | 14.4 | 71.3 | 10.1 | 16.3 | 14.9 | 47.7 | 7.0 | 10.0 | 7.5 | 10.5 | 4.6 |
| IL-7RA | 10.3 | 9.8 | 3.2 | 18.3 | 10.0 | 15.1 | 9.1 | 8.1 | 15.7 | 12.2 | 5.8 | 13.9 | 10.2 | 10.0 | 8.3 |
| IL-9 | 16.7 | 3.2 | 5.3 | 3.5 | 5.0 | 15.5 | 1.0 | 6.3 | 5.2 | 12.3 | 1.0 | 5.5 | 3.9 | 5.1 | 1.0 |
| IP-10/CXCL10 | 43.4 | 40.1 | 32.8 | 79.7 | 71.2 | 68.1 | 75.7 | 73.3 | 63.9 | 78.6 | 119.4 | 93.0 | 80.0 | 100.1 | 99.5 |
| LEPTIN | 56.5 | 39.5 | 35.1 | 12.7 | 6.3 | 98.7 | 16.8 | 48.4 | 18.4 | 76.7 | 39.5 | 23.9 | 38.3 | 29.0 | 11.3 |
| LIF | 3.5 | 4.5 | 6.3 | 8.6 | 6.4 | 110.1 | 5.7 | 6.6 | 6.7 | 5.6 | 5.1 | 1.0 | 5.8 | 4.7 | 4.7 |
| M-CSF | 1.0 | 1.0 | 1.0 | 1.0 | 1.0 | 1.0 | 1.0 | 1.0 | 1.0 | 1.0 | 1.0 | 1.0 | 1.0 | 1.0 | 1.0 |
| MCP-1/CCL2 | 1.0 | 1.0 | 1.0 | 14.7 | 8.2 | 11.2 | 3.5 | 1.0 | 3.3 | 1.0 | 5.9 | 1.0 | 1.0 | 1.0 | 1.0 |
| MCP-3/CCL7 | 59.0 | 85.3 | 113.6 | 353.4 | 429.7 | 285.3 | 327.8 | 140.7 | 227.1 | 369.9 | 135.2 | 67.9 | 58.5 | 116.6 | 92.0 |
| MIP-1a/CCL3 | 10.4 | 0.8 | 1.9 | 1.4 | 1.0 | 14.5 | 1.0 | 3.8 | 2.9 | 11.0 | 1.8 | 3.4 | 1.9 | 3.1 | 1.0 |
| MIP-1b/CCL4 | 5.7 | 5.5 | 4.0 | 39.6 | 28.9 | 21.3 | 20.9 | 15.0 | 17.4 | 32.6 | 33.2 | 21.8 | 13.1 | 14.9 | 16.6 |
| MIP-2 | 4.7 | 5.6 | 4.5 | 6.1 | 6.4 | 6.5 | 5.4 | 4.8 | 6.3 | 5.5 | 5.4 | 5.9 | 5.8 | 5.7 | 5.8 |
| RANTES/CCL5 | 26.8 | 10.1 | 12.8 | 10.5 | 13.3 | 26.9 | 13.2 | 13.9 | 13.2 | 21.2 | 14.4 | 14.7 | 12.2 | 14.5 | 13.8 |
| ST2/IL-33R | 1.0 | 1.0 | 1.0 | 1.0 | 1.0 | 1.0 | 1.0 | 1.0 | 1.0 | 1.0 | 1.0 | 1.0 | 1.0 | 1.0 | 1.0 |
| TNF-a | 1.0 | 1.0 | 1.0 | 3.4 | 3.7 | 3.7 | 1.0 | 1.0 | 1.0 | 1.0 | 1.0 | 1.0 | 1.0 | 1.0 | 1.0 |
| VEGF | 7.1 | 6.9 | 8.0 | 7.1 | 7.5 | 16.3 | 8.7 | 7.2 | 6.2 | 6.4 | 7.0 | 6.7 | 5.6 | 7.8 | 6.1 |
| sRANKL | 26.3 | 10.5 | 16.3 | 1.0 | 4.1 | 22.6 | 7.3 | 10.2 | 8.9 | 25.2 | 5.8 | 10.6 | 8.0 | 7.9 | 6.4 |

***Table S3.*** Functional enrichment. Pathways identified with gene set enrichment analysis (GSEA) using the Hallmark gene set.

| Sample | Pathway | FDR_Q_VAL | Norm.Enrichment |
|---|---|---|---|
| LNP-3 to SM102, Liver | MYC Targets V1 | 0 | -2.1898573 |
| | Unfolded Protein Response | 0 | -2.0541285 |
| | E2F Targets | 0.00127683 | -1.914668 |
| | Bile Acid Metabolism | 0.00072179 | 1.88673596 |
| | Interferon Gamma Response | 0.00095763 | -1.875752 |
| | Xenobiotic Metabolism | 0.00288716 | 1.79789733 |
| | TNFA Signaling VIA NFKB | 0.00280903 | -1.7886058 |
| | IL6 JAK STAT3 Signaling | 0.00297928 | -1.7798718 |
| | Interferon Alpha Response | 0.00711379 | -1.7127055 |
| | MTORC1 Signaling | 0.00941665 | -1.6832254 |
| LNP-5 to SM102, Liver | TNFA Signaling VIA NFKB | 0 | -2.364078 |
| | Interferon Alpha Response | 0 | 2.16394763 |
| | Myogenesis | 0.00404126 | -1.9371412 |
| | E2F Targets | 0.01154646 | -1.8123589 |
| | IL6 JAK STAT3 Signaling | 0.04041259 | -1.621349 |
| | Interferon Gamma Response | 0.03955367 | 1.62036116 |
| LNP-5 to SM102, Spleen | Interferon Alpha Response | 0 | 3.134800073 |
| | Interferon Gamma Response | 0 | 3.12053815 |
| | E2F Targets | 0 | -3.004694968 |
| | G2M Checkpoint | 0 | -2.748814384 |
| | Heme Metabolism | 0 | -2.627175439 |
| | Complement | 0 | 2.195932991 |
| | MYC Targets V2 | 0 | -2.175094241 |
| | Allograft Rejection | 0 | 2.136356098 |
| | MYC Targets V1 | 0 | -2.135753433 |
| | Inflammatory Response | 0 | 2.126372635 |
| | Coagulation | 0 | 2.106166588 |
| | TNFA Signaling VIA NFKB | 0.00014263 | 2.021970851 |
| | Myogenesis | 0 | -2.020599181 |
| | IL6 JAK STAT3 Signaling | 0.0001248 | 1.996375174 |
| | KRAS Signaling UP | 0.00022187 | 1.88409838 |
| | Mitotic Spindle | 0.00028678 | -1.844107054 |
| | Angiogenesis | 0.00059904 | 1.796195991 |
| | IL2 STAT5 Signaling | 0.00108917 | 1.754068203 |

***Table S4.*** Sequence of Fluc circRNA used in the study.

| | |
|---|---|
| IRES | TTTAAGTGTTGTGCCCAATCTCTTGACTCCTGCTGGAACCACCGACCAGTAGTGTCCAA<br>AATGCCAGGTGGAAAATCCTCCCTTCCCCTCTGGGCTTCATGCCCGGCATCCTCCCCCC<br>AGCCTGACGTGCCACAGGCTGTGCAAAGACCCCGCGAAAGCTGCCAAAAGTGGCAATT<br>GTGGGTCCCCCCTTTGTCAAGGCGTCGAGTCTTTCTCCCTTAAGGCTAGTCCTGTCAGTG<br>AACTCTGTCGGGCAACTAGTGACGCCACTGCATGCCTCCGACCTCGGCCGCGGAGTGCT<br>GCCCCCCAAGTCATGCCCCTGACCACAAGTTGTGCTGTCTGGCAAACATTGTCTGTGAG<br>AATGTTCCGCTGTGGCTGCCAAGCCTGGTAACAGGCTGCCCCAGTGTGCGTAATTCTCA<br>TCCAGACTTCGGTCTGGCAACTTGCTGTTAAGACATGGCGTAAGGGGCGTGTGCCAACG<br>CCCTGGAACGAGTGTCCACTCTAATACCCCGAGGAATGCTACGCAGGTACCCCTGGCTC<br>GCCAGGGATCTGAGCGTAGGCTAATTGTCTAAGGGTATTTTCATTTCCCACCCTCTTCTT<br>CTTGTTCATA |
| Optimized<br>Fluc<br>CDS | ATGGGGTCTGAAGATGCAAAGAATATCAAAAAGGGTCCCGCCCCTTTTTATCCTCTTGA<br>AGATGGCACCGCCGGCGAGCAGCTGCACAAAGCAATGAAAAGATACGCCCTGGTTCCT<br>GGAACTATTGCATTTACAGACGCTCACATCGAAGTTGATATTACTTATGCCGAATATTT<br>TGAAATGTCTGTTCGTCTGGCAGAAGCAATGAAAAGATACGGATTAAACACCAACCAT<br>CGGATCGTTGTATGCTCCGAGAACAGCCTGCAGTTTTTCATGCCTGTCCTGGGCGCCCT<br>GTTCATCGGCGTCGCTGTGGCCCCTGCAAACGACATCTACAACGAGAGAGAACTGCTG<br>AACAGCATGGGGATCTCCCAGCCTACTGTCGTCTTCGTCTCAAAGAAAGGATTACAGA<br>AAATCCTGAACGTACAGAAAAAGCTGCCCATCATCCAGAAAATCATCATCATGGATTC<br>TAAGACCGACTACCAGGGCTTCCAGTCTATGTATACCTTCGTCACCAGCCATCTGCCCC<br>CTGGATTTAATGAATATGATTTTGTCCCAGAATCTTTCGACAGAGATAAAACCATCGCC<br>CTGATCATGAACAGCTCTGGCTCCACAGGTTTACCTAAAGGAGTTGCTCTGCCCCATCG<br>GACCGCCTGTGTCCGGTTCAGCCATGCCCGGGACCCTATCTTCGGCAACCAGATTATCC<br>CTGATACCGCCATCCTGTCCGTTGTACCATTTCACCATGGTTTCGGCATGTTCACAACCC<br>TGGGCTATCTGATCTGCGGGTTTAGGGTCGTCCTGATGTATAGGTTCGAAGAAGAACTG<br>TTCCTGAGGTCACTGCAGGACTACAAGATCCAGTCTGCACTTCTGGTTCCTACTTTATTT<br>TCTTTCTTCGCAAAGTCCACACTGATCGACAAATACGACCTGTCCAATTTACATGAAAT<br>TGCATCCGGAGGAGCACCTCTTTCCAAGGAAGTTGGGGAAGCAGTTGCTAAGAGGTTC<br>CACCTGCCCGGAATTAGACAGGGCTATGGGCTGACCGAGACTACCAGCGCCATCCTGA<br>TCACCCCTGAGGGCGATGACAAACCTGGAGCAGTTGGGAAGGTGGTTCCTTTTTTCGAA<br>GCAAAGGTGGTTGATTTGGATACCGGCAAGACCCTGGGCGTCAATCAAAGAGGAGAAC<br>TGTGCGTCCGGGGCCCCATGATCATGTCTGGCTATGTAAATAATCCTGAGGCAACAAAC<br>GCCCTGATCGACAAAGATGGCTGGCTGCACAGCGGGGACATCGCCTACTGGGATGAAG<br>ATGAACATTTTTTCATCGTTGATCGTCTGAAGTCCCTGATCAAATACAAGGGTTATCAG<br>GTCGCTCCTGCAGAACTGGAGTCTATTTTACTGCAGCATCCTAATATCTTCGACGCTGG<br>TGTTGCTGGTTTACCTGATGACGATGCAGGTGAACTGCCCGCCGCTGTGGTTGTACTGG<br>AGCACGGGAAGACCATGACAGAAAAGGAAATTGTTGATTATGTAGCATCCCAGGTCAC<br>CACCGCCAAGAAACTGAGGGGTGGCGTCGTCTTCGTCGACGAGGTGCCTAAAGGATTA<br>ACAGGTAAACTGGATGCAAGAAAAATCCGCGAGATCCTGATCAAAGCAAAGAAAGGA<br>GGAAAAATCGCCGTTTAG |
